# A hidden mark of a troubled past: neuroimaging and transcriptomic analyses reveal interactive effects of maternal immune activation and adolescent THC exposure in the absence of overt behavioural disruptions

**DOI:** 10.1101/2024.08.05.606145

**Authors:** Mario Moreno-Fernández, Víctor Luján, Shishir Baliyan, Celia Poza, Roberto Capellán, Natalia de las Heras, Miguel Ángel Morcillo, Marta Oteo, Emilio Ambrosio, Marcos Ucha, Alejandro Higuera-Matas

## Abstract

Maternal exposure to infections during gestation has been shown to predispose individuals to the development of schizophrenia. Additionally, clinical data suggest that cannabis use could trigger the onset of schizophrenia in vulnerable individuals. However, the direction of causality remains unclear. To elucidate this issue, we utilised a rat model of maternal immune activation, combined with exposure to increasing doses of THC during adolescence, in both male and female rats. We investigated several behaviours in adulthood that resemble specific symptoms of schizophrenia, including impairments in working memory, deficits in sensorimotor gating, alterations in social behaviour, anhedonia, and potential changes in implicit learning (conditioned taste aversion). Furthermore, we conducted a longitudinal positron emission tomography (PET) study to identify affected brain regions and subsequently collected brain samples from one of these regions (the orbitofrontal cortex) for RNA sequencing analyses. These analyses were also performed on peripheral blood mononuclear cells to identify peripheral biomarkers. Notably, while no overt behavioural disruptions were observed, our PET scans revealed several brain alterations dependent on the combination of both exposures. Additionally, the transcriptomic studies demonstrated that maternal immune activation affected glutamatergic and serotoninergic genes, with the combination of both exposures shifting the expression from down-regulation to up-regulation. In peripheral cells, interactive effects were observed on inflammatory pathways, and some genes were proposed as biomarkers of the disease. These results suggest that the combination of these two vulnerability factors leaves a lasting mark on the body, potentially predisposing individuals to the disease even before behavioural alterations manifest.

## INTRODUCTION

At the end of the 19th century, the French psychiatrist Jacques-Joseph Moreau was among the first to describe psychotic-like experiences following the ingestion of high doses of cannabis (Moreau, 1973). Since then, a growing body of evidence has linked cannabis consumption with an increased risk of developing disorders within the schizophrenia spectrum (Groening et al., 2024; Hjorthøj et al., 2023; Petrilli et al., 2022). However, the available data do not allow for definitive conclusions regarding the direction of causality. Indeed, Mendelian randomisation studies even suggest a bidirectional association between cannabis use and psychosis (Groening et al., 2024). To address this conundrum, animal studies are essential. Numerous animal models have been developed to replicate various symptom dimensions of schizophrenia. Among these, models based on maternal immune activation (MIA) have been pivotal in studying susceptibility and resilience factors (Meyer, 2019). There is substantial evidence linking prenatal infections, including bacterial infections, with an increased risk of developing psychosis (Cheslack-Postava and Brown, 2022). Utilising these models, researchers have combined MIA, serving as a first hit, with cannabinoid exposure during adolescence, acting as a second, triggering hit, to examine the behavioural and neural sequelae of each hit and, importantly, their potential synergistic effects in the adult offspring (see Martín-Cuevas et al., (2023) for a review).

In previous work, we demonstrated that LPS-induced MIA and chronic, low-dose adolescent Δ9-tetrahydrocannabinol (THC) exposure (ATE) did not synergistically alter several behavioural dimensions relevant to schizophrenia, such as working memory, pre-pulse inhibition, social interaction, sucrose preference, or incidental association formation (Moreno-Fernández et al., 2024). These findings align with those of other laboratories, which also reported a lack of interaction between MIA (induced by the viral RNA analogue Poly I:C) and ATE in behavioural tests of emotional (open field, elevated plus maze, forced swimming test) and cognitive processes (Morris water maze, prepulse inhibition, and fear conditioning) (Stollenwerk and Hillard, 2021). In a more recent study, Guma and colleagues also observed a lack of synergy between MIA and ATE in behavioural tests pertinent to schizophrenia (open field, prepulse inhibition, and social interaction in the three-chamber test). However, when they examined brain structure using magnetic resonance imaging on postnatal day 50 (five days after the last THC injection), they noted significant effects of the combination of both hits on the volume of the nucleus accumbens, striatum, somatosensory, anterior cingulate, and entorhinal cortices, as well as the subiculum. Nevertheless, these changes were transient and had dissipated by postnatal day 80 (Guma et al., 2023). Other studies have even suggested protective effects of ATE (Lamanna-Rama et al., 2024).

These data are intriguing but are based on THC administration protocols with either a low (3 mg/kg i.p. in rats on alternate days) (Moreno-Fernández et al., 2024), medium (5 mg/kg i.p. in mice, daily injections) (Guma et al., 2023) or high dose (10 mg/kg i.p. daily injections) (Lamanna-Rama et al., 2024); or oral administration in cereal pieces (3 mg/kg in mice) (Stollenwerk and Hillard, 2021) which complicates the assessment of the actual THC exposure levels in the animals. It may be more informative to utilise a model of ATE with increasing doses, which could better reflect the pattern of problematic cannabis use associated with the development of psychotic disorders in humans (Groening et al., 2024).

In light of this evidence, the present study combines a model of LPS-induced MIA in rats with ATE using increasing doses, to investigate behavioural alterations relevant to schizophrenia in male and female offspring. Crucially, given that prior experiments indicated that brain alterations might precede overt behavioural impairments, we employed positron emission tomography (PET) to obtain data on brain function, complementing these findings with RNA sequencing (RNAseq) analyses to gain deeper insights into the gene networks involved. Furthermore, we extended these analyses to peripheral blood mononuclear cells (PBMCs) to determine whether changes observed in the brain could also be detected peripherally, thus identifying potential biomarkers of the disease. Additionally, adopting a preventive approach, we explored predictive variables in the dams (behavioural traits and physiological responses to the immune challenge) that might indicate the severity of symptoms in the offspring.

## MATERIALS AND METHODS

For a full description of the methodology and materials used, please refer to the supplementary materials and methods.

Figure S1 provides a schematic diagram of the experimental timeline.

### Animals

All experiments were approved by the Ethics Committee of UNED and the Autonomous Community of Madrid (PROEX 177.4/20). The animals were maintained and treated in accordance with the European Union Laboratory Animal Care Guidelines (EU Directive 2010/63/EU), with meticulous adherence to the “Principles of Animal Welfare.” All animals were Sprague-Dawley (SD) CD rats obtained from Charles River.

### Mating and LPS Administration

Females were mated with males at a 1:1 ratio. On gestational days (GD) 15 and 16, the rats received intraperitoneal injections of 50 µg/kg lipopolysaccharides (LPS) dissolved in sterile saline solution (0.9% NaCl solution; Braun) (n=11) or saline (n=12). Rectal temperatures were recorded immediately before the injections and 60, 120, and 240 minutes afterward to indicate MIA. Body weight was measured on the days of injection and on two additional consecutive days (from GD15 to GD18). All pups were weaned on postnatal day (PD) 22 and grouped according to sex and prenatal exposure. They were then assigned to either the THC or vehicle group, ensuring each group included pups from all dams.

### Adolescent THC Exposure

THC was obtained from THCPharm (Frankfurt, Germany). Vehicle or increasing doses of THC were administered intraperitoneally at a volume of 2 mL/kg. From PD30 to PD32, a dose of 2.5 mg/kg was administered; from PD33 to PD36, the dose was increased to 5 mg/kg; and from PD37 to PD40, the dose was 10 mg/kg (protocol adapted from Rubino et al., 2008).

### Study of potential early markers in the dams predictive of behavioural deficits in the offspring

#### Elevated plus maze

Anxiety-like behaviour was assessed approximately one week before mating, at the age of 10–12 weeks, using an elevated plus maze. The test duration was 5 minutes, following the method described by Higuera-Matas et al., (2009).

#### Locomotor activity

Two days after the elevated plus maze test, locomotor activity in a novel environment was analysed using an open field maze. Rats were placed in the centre of the maze and allowed to explore for 40 minutes.

#### Immune Reactivity and Flow Cytometry Assay

The immune response magnitude to additional LPS injections was analysed in all dams approximately two weeks after weaning. On the first day, half of each group received an LPS injection (50 µg/kg, i.p.), while the other half received a saline solution. Following a washout day, the procedure was repeated, injecting LPS or saline into those who had not received it previously. Monocyte, B lymphocyte, and T lymphocyte populations in PBMC were analysed via flow cytometry using a blood sample obtained from the tail vein.

#### Behavioural Tests in the Offspring

Upon reaching adulthood, a comprehensive set of behavioural tests was conducted to evaluate behaviours analogous to the different symptoms of schizophrenia. Cognitive alterations were assessed in the working memory (Y maze) and sensorimotor gating (prepulse inhibition of the acoustic startle response) domains. For negative symptoms, social interactions and sucrose preference (a measure of anhedonia) were analysed. Conditioned taste aversion was used to assess implicit learning alterations.

#### Working Memory

The Y-maze was employed to assess working memory, with the percentage of alternation calculated as the ratio of the number of triads (three consecutive entries into different arms) to the maximum possible number of triads (total entries minus two) (Conrad et al., 1997; Lainiola et al., 2014; Prieur and Jadavji, 2019).

#### Social Interaction

This study focused on sociability and social novelty preference. A conventional three-chamber arena was used to study these variables. In phase 1 (8 minutes exploration), one cylinder contained a conspecific rat of the same sex (Conspecific 1), while the other contained an inanimate object (a stuffed rat). In phase 2, a new conspecific rat (Conspecific 2) replaced the object, and the behaviour of the test rat was recorded for another 8 minutes. Interaction time was measured as the time the rat’s head was within a 5 cm radius of the cylinder and facing it.

#### Test for Incidental Associations and Conditioned Taste Aversion

Animals were provided with 1 hour of water access in the morning, with scents and/or flavours dissolved in the drinking water. They were single-housed during this test. Following a drinking training phase, animals underwent six sessions of compound exposure (Sensory Preconditioning Phase), each consisting of two days of 1-hour exposure to one of two odour-taste combinations: 0.01% almond odour with 5 mM NaCl taste, and 0.05% banana odour with 0.316 mM HCl. The conditioning phase involved exposing all groups to one of the odours for 1 hour, followed by an injection of LiCl or saline. This cycle was repeated three times. The test phase included a mediated test to assess aversion to flavours preconditioned with LiCl-paired odours and a direct test to measure aversion to the LiCl-paired odours.

#### Sensorimotor Gating

Prepulse inhibition of the startle response was used as a measure of sensorimotor gating. Prepulses were set at 4 or 12 decibels (dB) above background noise (65 dB) with intervals of 30 or 120 milliseconds (ms) between prepulse and pulse (120 dB).

#### Anhedonia

The anhedonia test involved four daily sessions of 1-hour exposure to two water bottles, one containing tap water and the other 1% sucrose solution.

### Positron Emission Tomography

The effects of MIA, ATE, and their interaction on [18F]-2-fluoro-2-deoxy-D-glucose (18F-FDG) brain uptake were studied in a new subset of 80 Sprague-Dawley rats (40 males and 40 females), divided into four groups (n=10) at two time points: PND 60 (PET1) and PND 120 (PET2). PET-CT studies were conducted using a small-animal SuperArgus PET/CT (SEDECAL, Madrid, Spain). Statistical analysis was performed using SPM software (http://www.fil.ion.ucl.ac.uk/spm/software/spm12/). A two-way ANOVA (p < 0.01 uncorrected) was used to assess the effects of MIA, ATE, and their interaction on 18F-FDG uptake in the brain, for each sex and time point. To minimise type I errors, only clusters with p < 0.05 (uncorrected) were considered.

### Massive Parallel RNA Sequencing

Massive parallel RNA sequencing was performed on another subset of rats (n=3 per group) at the Genomics Unit of the Madrid Science Park, using an Illumina® NovaSeqTM 6000 instrument with the NovaSeqTM Reagent 1.5 kit (100 cycles) in a single-read 1×100 run. Sequences were mapped using the TopHat program within the G-PRO suite (Biotechvana) and a differential expression analysis was conducted using the CuffDiff program. Gene expression data are available in the Gene Expression Omnibus (GEO) repository.

### Data Analysis

#### Video Analysis

Animal behaviour was analysed using the ANY-maze Video Tracking System v.6.32 (Stoelting Co.) or higher.

#### Indices and Correlations

To investigate early markers predictive of deficits, a predictive index for the dams (DI) was calculated using normalised data from various measures, including elevated plus maze performance, locomotor activity, rectal temperature variation after LPS administration, weight loss, and T lymphocyte percentage change post-weaning. Similarly, an offspring behavioural index (OI) was created by normalising and averaging each litter’s test results. Correlation analyses were performed between the indices and/or their components.

#### Data Normalisation and Statistics

Refer to the Supplementary Materials and Methods for detailed information.

## RESULTS

### MIA induction

LPS administration induced a hypothermic response on GD15, evidenced by a significant Time*MIA interaction (F(2, 27) = 16.28, p < 0.001, η^2^p = 0.538) (Figure S2A and B). This effect was absent on GD16, suggesting potential habituation to the hypothermic effects of endotoxin exposure. LPS rats also exhibited reduced bodyweight gain compared to their weight on GD15 (Figure S2C and Table S2).

### Behavioural Screening

#### Working Memory in the Y Maze

Analyses of the percentage of alternations in the offspring, assessing working memory, revealed a significant Sex effect (F(1, 77) = 7.66, p = 0.007, η²p = 0.090), indicating better performance in males. No effects due to MIA were observed, nor were there significant differences caused by ATE or its interaction with prenatal LPS exposure (Figure S3A and Table S3).

#### Social Interaction and Social Novelty Preference

A repeated measures ANOVA indicated that animals spent more time interacting with the conspecific than with the inanimate object (F(1, 75) = 51.98; p = 0.000, η^2^p = 0.409). No significant effect of LPS-induced MIA on preference percentage was found, nor any effect due to ATE (Figures 3B and Table S4).

Regarding preference for social novelty, a general preference for the newly introduced rat was observed, as analysed by a repeated measures ANOVA of interaction times (F(1, 79) = 20.91; p = 0.000, η²p = 0.209). An interaction between MIA and ATE was identified when analysing the percentage of preference for the new rat (Table S4). Simple effects analysis revealed that ATE increased preference for social novelty in animals exposed to LPS during gestation (Figure S3C). This effect was not observed in animals exposed to saline during gestation (Table S4).

#### Prepulse Inhibition

Analyses of the prepulse inhibition test revealed no effects due to MIA, ATE, or their interaction under any condition (Table S5). However, greater response inhibition was observed in females compared to males in the 4dB_30ms condition (Figure S3D). Additionally, a trend towards a significant Sex*MIA interaction (p = 0.055) was noted in the habituation data, suggesting that MIA may reduce habituation in males (Figure S3E).

#### Sucrose Preference

The sucrose preference test aimed to identify potential deficits analogous to anhedonia observed in humans with schizophrenia. All animals showed a general preference for the sucrose solution (F(1, 68) = 481.78; p = 0.000, η²p = 0.876). No differences in preference percentages due to MIA, ATE, or their interaction were observed (Figure S3F), nor were there any differences due to sex.

#### Conditioned Taste Aversion

A clear effect of LiCl was demonstrated by significantly lower consumption of the conditioned odour solution in all groups (F(1, 80) = 322.83; p = 0.000, η²p = 0.801) (Figure S3G). A significant CS+*Sex effect was noted: while both males and females exhibited conditioned taste aversion, the effect size was larger in males (η²p = 0.884) than in females (η²p = 0.650). No other significant effects were found.

### Early Predictive Markers of Behavioural Disruption in the Offspring

Potential early markers in the dams that could predict the severity of deficits in the offspring were explored. Correlations between the Offspring Index (OI), the Dam Index (DI), and their components were analysed. A positive correlation was found between the OI and the change in the dam’s weight after the first LPS injection (Table S6). This result suggests that greater weight loss following immunogen exposure during gestation correlates with poorer performance in the behavioural tests.

### Brain Activity Assessment by PET

Our longitudinal analysis of the interactions between MIA and ATE at two time points (PND60 - T1 - and PND120 - T2) on brain activity revealed distinct patterns and significant sex differences (Figure 1). One pattern showed increased activation in animals experiencing both MIA and ATE (positive interaction) compared to those exposed only to MIA. The second pattern indicated decreased activation in animals subjected to both MIA and ATE (negative interaction) relative to MIA rats alone.

**Figure 1.**
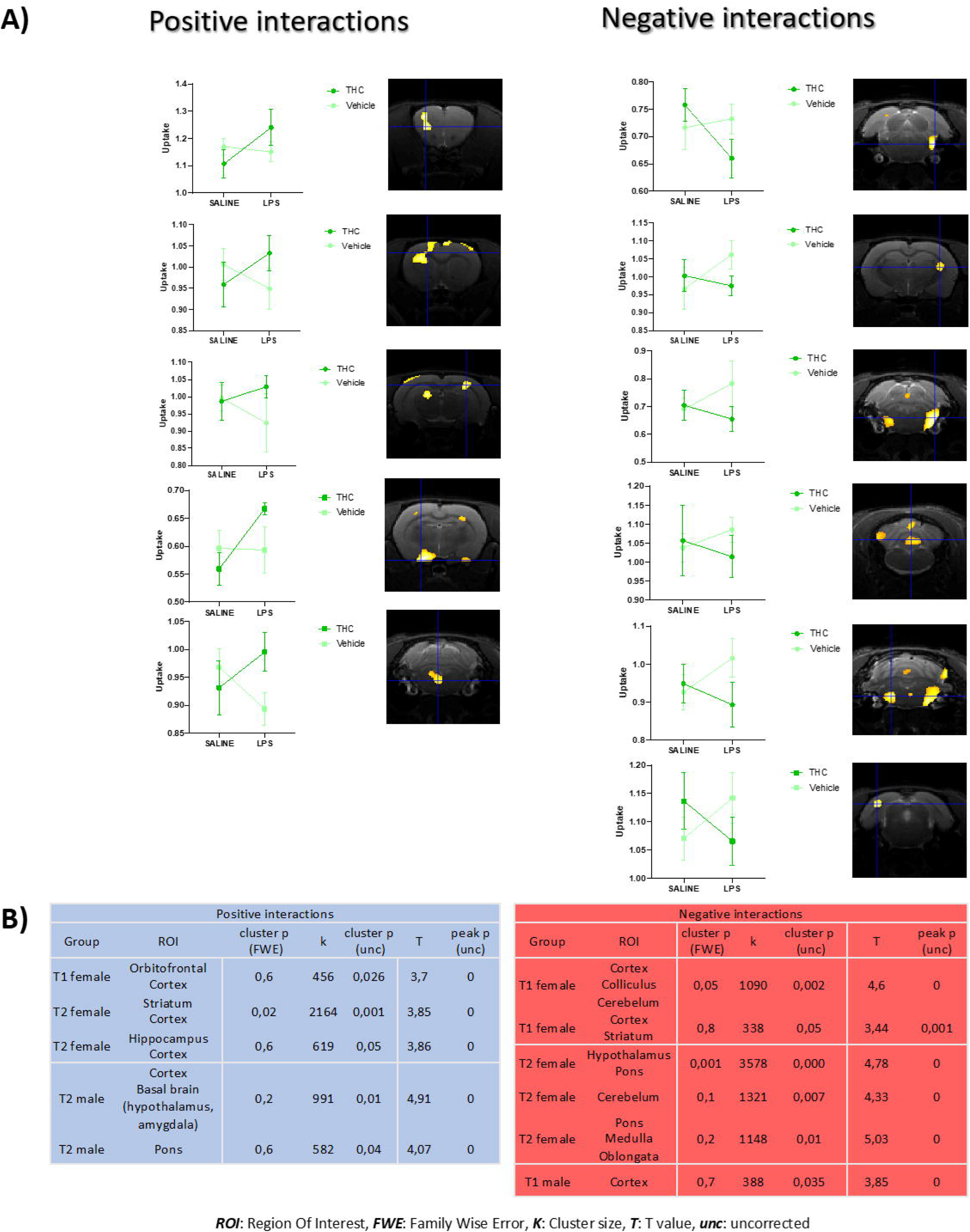
Interactive effects on brain activity assessed by PET in males and females. A) Study of the interactions (positive in blue and negative in red) of MIA and ATE for each time point (T1 and T2) and sex (female in purple and male in brown) that were above a threshold of p value of 0.01 uncorrected for the voxel and a p value < 0.05 uncorrected for the whole cluster, to avoid type I error. For each type of interaction, plots of the uptake effect and affected regions overlaid on an MRI template are shown. B) Detailed SPM12 data per type of interaction, positive in blue and negative in red, with the statistical data of each cluster.

At T1, females exposed to MIA and ATE exhibited increased activation in clusters including the lateral orbitofrontal cortex (OFC) and motor cortex. At T2, this interaction disappeared in the OFC but emerged in the dorsolateral striatum, hippocampus, and parts of the thalamus and visual cortex. In males, at T2 we detected a positive interaction pattern in the amygdala, lateral hypothalamus, and central pons.

Regarding negative interactions, small clusters in the lateral pons and striatum were noted at T1, with most effects occurring in the caudal brain regions, especially the cerebellum and lateral pons. There was also a small cluster of interactions in the visual cortex of males at T1. Notably, a sex-specific effect was observed: males showed a positive interaction in the pons at PND 120, while females showed a negative interaction. See Figures S4 and S5 for a detailed description of the main effects of MIA and ATE alone.

### Orbitofrontal and Peripheral Mononuclear Cell Transcriptomes

#### Orbitofrontal Transcriptome

Given the involvement of the orbitofrontal cortex (OFC) in various aspects of schizophrenia and the effects observed in our PET analysis, we conducted a transcriptomic analysis of the OFC.

In males, differential expression analysis revealed 1,737 differentially expressed genes (DEGs) (p < 0.05 adjusted) due to MIA, the majority of which were downregulated (Figure S6). The impact of ATE was notably smaller, affecting the expression of only 168 genes. Interestingly, the combination of MIA and ATE did not significantly increase the number of DEGs compared to control animals, with most genes being upregulated when exposed to both insults. The distribution of genes indicated that MIA induced the largest set of specific gene expression alterations (79.4% of all DEGs in the male study) (Figure S7A). Notably, 6.6% of DEGs were shared between MIA and ATE, indicating interactive effects.

Further analysis on how latent gene expression alterations induced by MIA could be modified by ATE (LPS+THC vs. LPS+VEH comparison) revealed a significant synergistic effect, with 2,620 DEGs identified. In contrast, the number of DEGs induced by ATE as a function of the MIA background (SAL+THC vs. LPS+THC comparison) was much smaller (270 DEGs) (Figure S6).

In females, MIA had a substantial effect, altering the expression of 1,185 genes, similar to the effect observed in males (Figure S6). The impact of ATE was considerably smaller than that of MIA, with changes in the opposite direction, as most genes were upregulated. The combination of both factors resulted in an intermediate effect (224 DEGs), significantly lower than MIA alone (1,185 DEGs) but slightly higher than ATE alone (136 DEGs).

Further analysis of how the latent effects of MIA could be modified by ATE in females showed a less pronounced effect on the DEG profile compared to males. The effect of ATE in MIA animals did not differ significantly from controls as observed in males. When comparing the comparisons indicative of interactive effects (LPS+VEH vs. LPS+THC and SAL+THC vs. LPS+THC), we found that 11.4% of the total affected genes were common to both (Figure S7B).

Following this quantitative analysis, we conducted a gene ontology enrichment analysis. In males, MIA affected ontologies related to neuron projection development and its regulation, synaptic signalling and organisation, and behaviour (Figure S8B). The general trend of downregulation included genes involved in the glutamatergic and serotoninergic systems, such as Gria1, Gria2, Gria3, Grin2a, Htr2a, and Htr2c (Figure S8A). Hierarchical cluster analysis of the main five ontologies (‘neuron projection development’, ‘synaptic signalling’, ‘regulation of neuron projection development’, ‘synapse organisation’, and ‘behaviour’) revealed two main clusters. One consisted of upregulated genes, with two sub-clusters (one with clearly upregulated genes like Npas4, Arc, Fos, Egr1, and Egr2; and another with a mixed pattern including Vgf and Bmp4). The other cluster comprised downregulated genes, with sub-clusters including a mixed pattern such as Cdnf and Epha3 and a clearer upregulation such as Grin2a and Cacnb4 (Figure S8D).

A similar pattern emerged in females, with the top five ontologies being ‘modulation of chemical synaptic transmission’, ‘synapse organisation’, ‘behaviour’, and ‘neuron projection development’ (Figure S9B). Notably, the highlighted genes from the volcano plot in males, except Grm2, were all downregulated in females as well. The top five ontologies revealed two clusters, indicating the previously mentioned upregulation and downregulation, predominantly in MCS and RNPD, respectively (Figure S9D).

We then examined whether ATE could exacerbate or alter the gene expression patterns induced by MIA in the OFC. In males, a general shift from downregulation to upregulation was observed in synapse-related genes (Figure 2A). Significant ontologies affected included ‘synaptic signalling’, ‘synapse organisation’, and ‘modulation of chemical synaptic transmission’. Importantly, these genes were not affected by ATE alone, as shown by the SAL+VEH vs. SAL+THC comparison results (Figure S10A).

**Figure 2:**
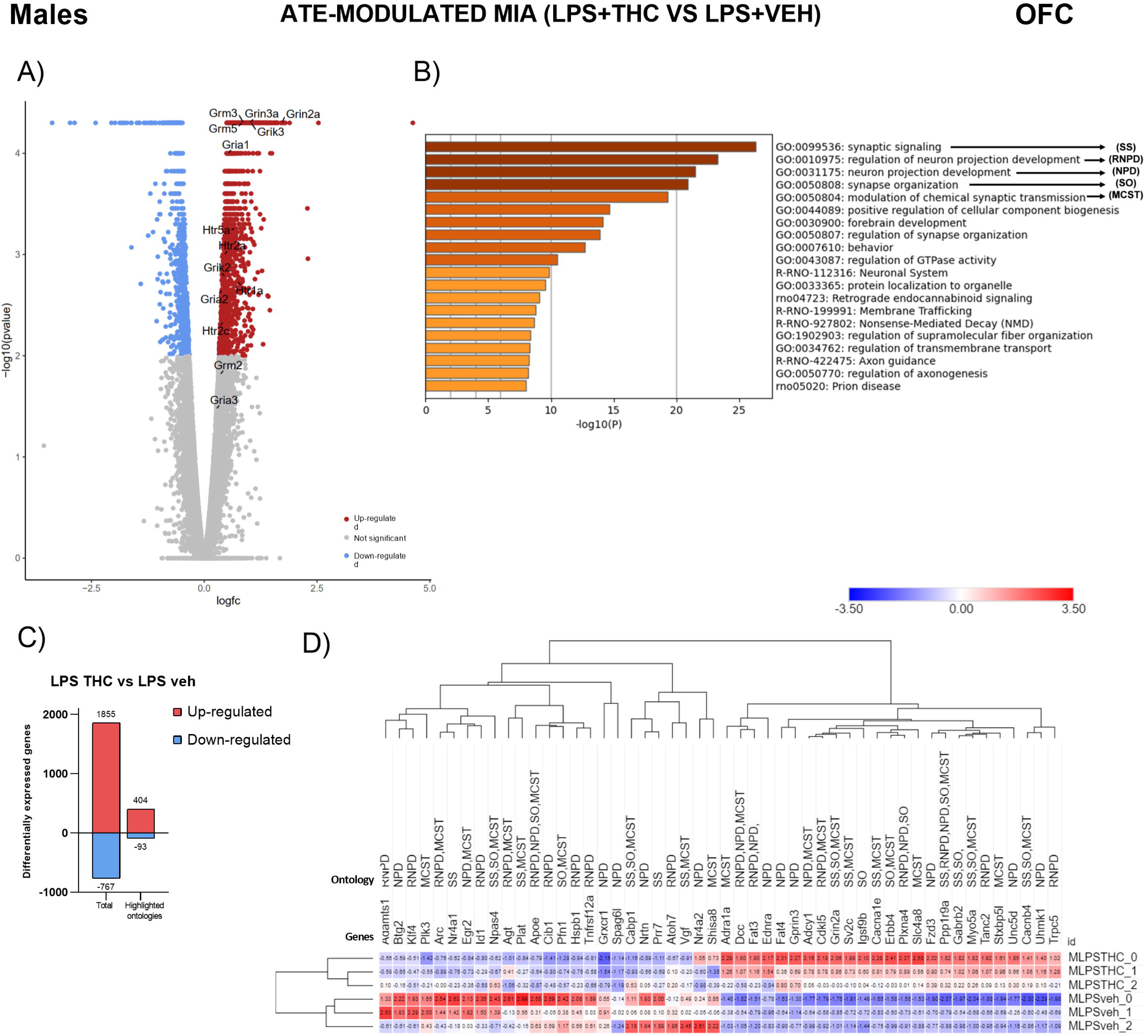
Modulation of MIA effects by ATE on OFC transcriptome in males. A) −Log10 of the unadjusted p value is plotted against the logarithm of FC (fold change). Genes differentially expressed upregulated are shown in red, while those downregulated are shown in blue (adjusted p value < 0.05). Genes of glutamatergic and serotoninergic receptors, or their subunits, affected by exposure to LPS, THC, or their combination are depicted. B) Ontologies significantly enriched with differentially expressed genes. C) Graphical representation of the number of differentially expressed genes, both total (Total) and only from the 5 highlighted ontologies (highlighted ontologies). D) Heatmap showing the number of fragments from each sample normalized to Z-score using the mean and standard deviation of fragments from all males. All genes belonging to the highlighted ontologies are represented.

A similar pattern emerged in the females, albeit to a much lesser extent (Figure 3). Notably, Grin2a was one of the genes exhibiting this shift in expression in both males and females (see Discussion). On its own, ATE regulated genes associated with myelination and plasticity ontologies (Figure S11).

**Figure 3:**
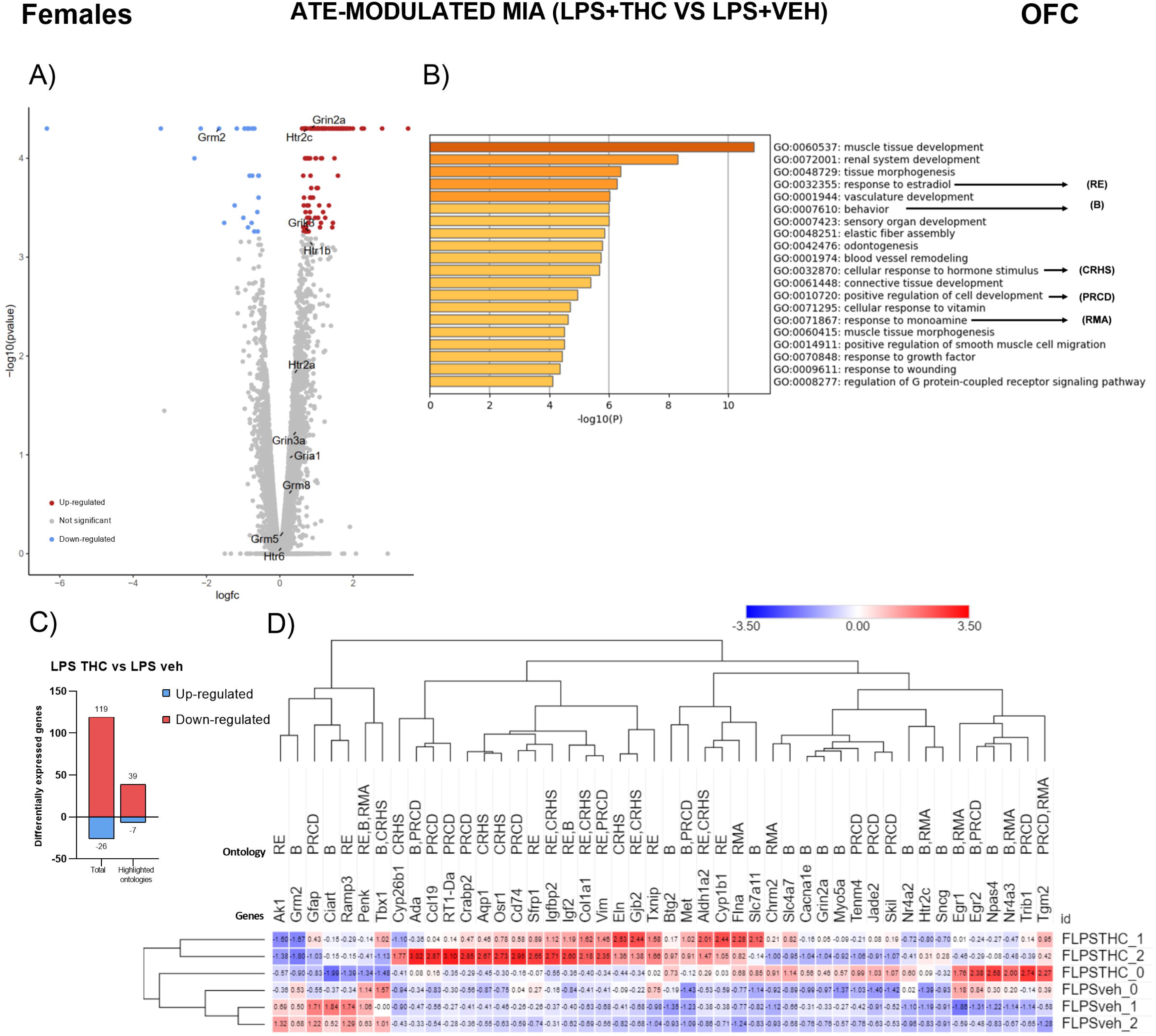
Modulation of MIA effects by ATE on OFC transcriptome in females. A) −Log10 of the unadjusted p value is plotted against the logarithm of FC (fold change). Genes differentially expressed upregulated are shown in red, while those downregulated are shown in blue (adjusted p value < 0.05). Genes of glutamatergic and serotoninergic receptors, or their subunits, affected by exposure to LPS, THC, or their combination are depicted. B) Ontologies significantly enriched with differentially expressed genes. C) Graphical representation of the number of differentially expressed genes, both total (Total) and only from the 5 highlighted ontologies (highlighted ontologies). D) Heatmap showing the number of fragments from each sample normalized to Z-score using the mean and standard deviation of fragments from all males. All genes belonging to the highlighted ontologies are represented.

These gene expression regulation patterns are particularly relevant and may provide a potential disease signature. To determine if any of these changes could be detected peripherally, we conducted an RNAseq study on the PBMCs of these animals.

#### PBMC transcriptome

Differential expression analysis in males revealed a total of 388 DEGs (p < 0.05 adjusted) due to MIA, with most being downregulated (Figure S12). Similarly, ATE induced a higher number of DEGs (603), also predominantly downregulated. The combination of both factors increased the total number of DEGs to 687, while still maintaining a majority of downregulated genes. In terms of DEG distribution, ATE showed a greater influence than MIA, with only 5.7% of genes being exclusively altered by LPS-induced MIA, compared to 23.2% by ATE (Figure S13A). Additionally, 26.4% of genes were exclusively differentially expressed in the presence of both factors.

When examining how latent gene expression changes induced by MIA could be modified by ATE (LPS+THC vs. LPS+VEH comparison), a lower number of DEGs (110) was observed, with a roughly equal proportion of genes being up- or down-regulated. Similarly, the number of DEGs induced by ATE as a function of the MIA background (SAL+THC vs. LPS+THC comparison) was found to be 121, fewer than those due to the effects of either factor alone or their combination. Regarding distribution, when comparing each impact with the combination, MIA altered 37% of the genes, ATE 42.7%, and the remaining 20.3% were shared between both comparisons (Figure S13A).

In females, MIA appeared to have a weaker effect (only 15 DEGs) compared to males (Figure S12). ATE was associated with a higher number of DEGs (148), but unlike in males, most of these genes were upregulated. The combination of both factors synergistically increased transcriptomic alterations to 319 DEGs. In this case, 61% of the genes were exclusively altered by the combined impact, confirming the previously mentioned synergistic effect (Figure S13B).

In studying how latent gene expression changes induced by MIA could be modified by ATE (LPS+THC vs. LPS+VEH comparison), a synergistic effect was again observed, with 252 DEGs, most of which were upregulated. We then examined the effects of ATE depending on the MIA background (SAL+THC vs. LPS+THC comparison) and found a lower number of DEGs (107), with the majority also being upregulated (Figure S12). These results indicate significant sex-dependent differences, such as the general direction of differential expression, the weaker effects in females, and the absence of synergistic effects in males.

Following this quantitative analysis, we conducted a gene ontology enrichment analysis. In males, MIA was associated with a significant enrichment in the ‘response to wounding’, ‘cell activation’, and ‘haematopoiesis’ ontologies (Figure S14B). We highlighted several genes important to these ontologies, such as S100a8 and S100a9 (downregulated), and genes involved in chemokine signalling (Cxcl2 and Cxcl10), cytokines (Il1b), or toll-like receptors (Tlr7, Tlr8, and Tlr13), which were upregulated (Figure S14A). Hierarchical cluster analysis of DEGs within these highlighted ontologies identified two main clusters, corresponding to up- and downregulated genes. Among the upregulated genes, two sub-clusters were found: one with strong upregulation (including Ccrl2, Cxcl2, Icam1, Nfkbiz, Cd59, Il1b, Slc4a1, and Spta1) and another with a mixed pattern (including Tlr7, Tlr13, or Cd180). Among the downregulated genes, one cluster comprised S1008 and S1009, while another included genes involved in the regulation of cell activation.

In females, the effect of MIA was much more limited, with only 15 DEGs. Notably, two genes upregulated in males, Il1b and Cxcl2, were downregulated in females (Figure S15A).

We then examined how the latent effects of MIA could be modified by ATE. In males, ATE shifted the MIA-associated upregulation to downregulation of certain genes (such as Cxcl2 and Il1b) (Figure 4A) and prevented the upregulation of other MIA-affected genes (such as the toll-like receptor genes). Additionally, the S100a8 and S100a9 genes, downregulated by MIA, were upregulated by ATE. The top three affected ontologies were ‘inflammatory response’, ‘chronic inflammatory response’, and ‘IL-17 signalling pathway’ (Figure 4B). Some of these genes (such as S100a8 and S100a9) were affected by ATE alone, in the same direction as MIA. Interestingly, ATE also influenced genes of the toll-like receptor family (Tlr7, Tlr8, and Tlr13) (Figure S16A).

**Figure 4:**
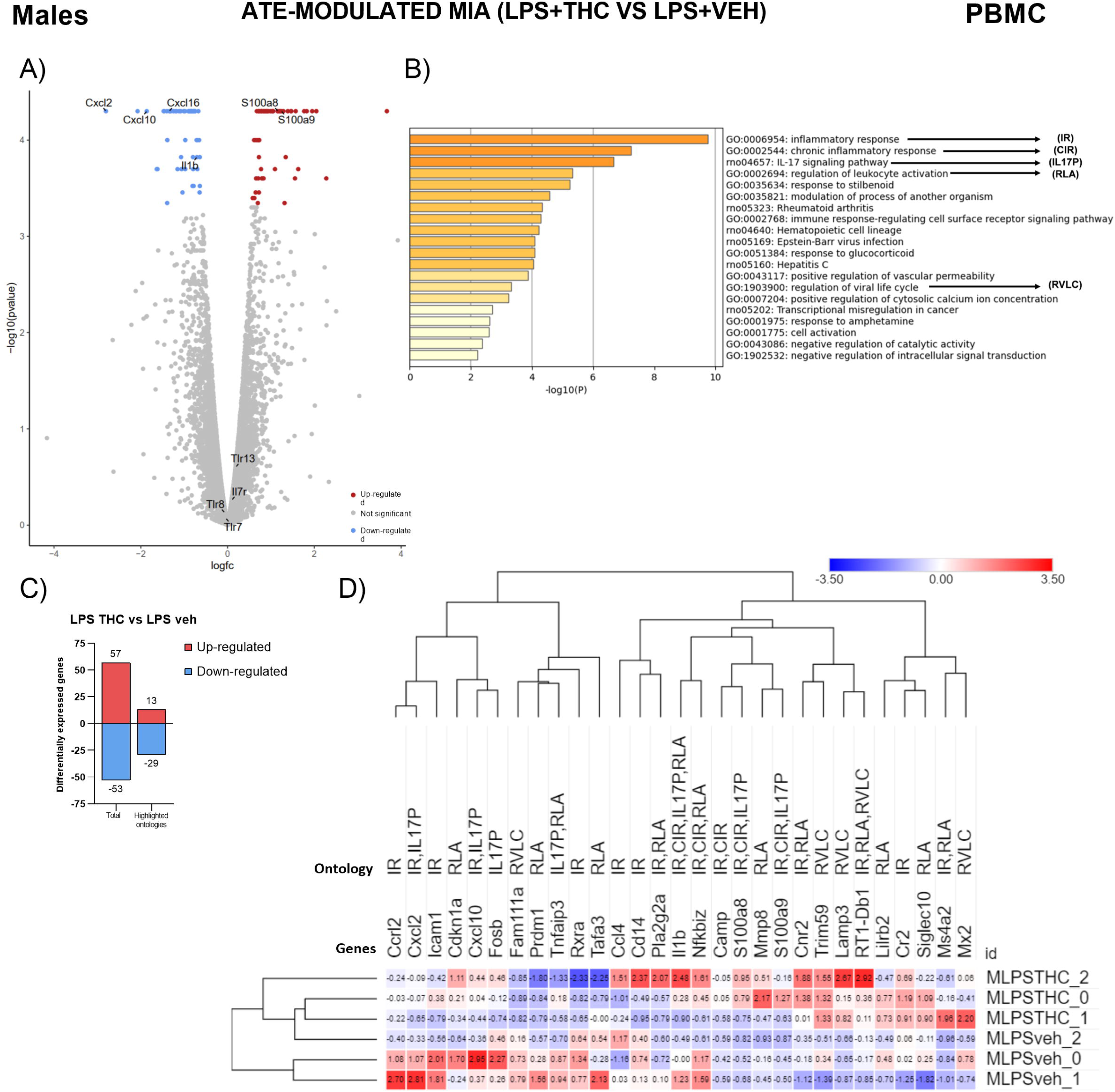
Modulation of MIA effects by ATE on PBMCs transcriptome in males. A) −Log10 of the unadjusted p-value is plotted against the logarithm of FC (fold change). Genes differentially expressed upregulated are shown in red, while those downregulated are shown in blue (adjusted p-value < 0.05). Genes of glutamatergic and serotoninergic receptors, or their subunits, affected by exposure to LPS, THC, or their combination are depicted. B) Ontologies significantly enriched with differentially expressed genes. C) Graphical representation of the number of differentially expressed genes, both total (Total) and only from the 5 highlighted ontologies (highlighted ontologies). D) Heatmap showing the number of fragments from each sample normalized to Z-score using the mean and standard deviation of fragments from all males. All genes belonging to the highlighted ontologies are represented.

In females, ATE modulated MIA effects, increasing the number of DEGs compared to the MIA comparison. The Il1b gene shifted expression direction from downregulation to upregulation, and some genes not significantly regulated by MIA were upregulated (such as S100a8 and S100a10) (Figure 5A). Notably, the most significantly enriched gene ontologies were related to erythrocyte physiology (Figure 5B).

**Figure 5:**
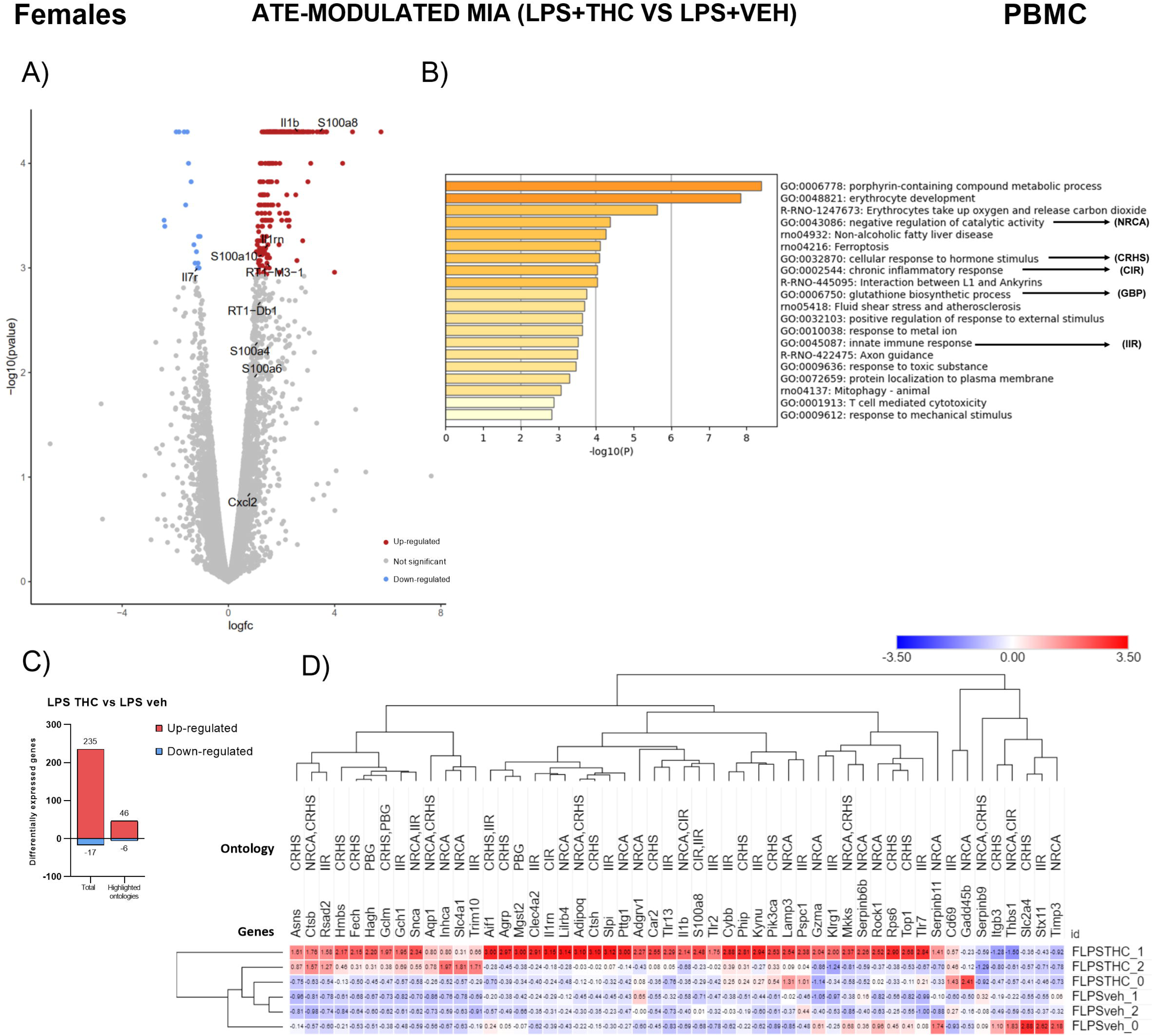
Modulation of MIA effects by ATE on PBMC transcriptome in females. A) −Log10 of the unadjusted p-value is plotted against the logarithm of FC (fold change). Genes differentially expressed upregulated are shown in red, while those downregulated are shown in blue (adjusted p-value < 0.05). Genes of glutamatergic and serotoninergic receptors, or their subunits, affected by exposure to LPS, THC, or their combination are depicted. B) Ontologies significantly enriched with differentially expressed genes. C) Graphical representation of the number of differentially expressed genes, both total (Total) and only from the 5 highlighted ontologies (highlighted ontologies). D) Heatmap showing the number of fragments from each sample normalized to Z-score using the mean and standard deviation of fragments from all males. All genes belonging to the highlighted ontologies are represented.

ATE also upregulated the S100a8 gene (Figure S17A) in females, but the other genes modulated in the comparisons were not significantly altered. The most represented ontology among DEGs in this comparison was also related to erythrocyte physiology (Figure S17B).

#### Search for common DEGs in the OFC and PBMC transcriptomes

A significant aim of this study was to identify DEGs in the OFC that were also regulated in the PBMC transcriptome, with the goal of discovering potential peripheral biomarkers of the combined vulnerability induced by MIA and ATE. In males, 49 genes were modulated by MIA in both the OFC and PBMCs. Of these, 12 were modulated in the same direction. Notably, Pde5a and Pde4d showed some of the highest fold-changes in PBMCs, facilitating their detection in blood samples, with similar or higher fold-changes in the OFC. Interestingly, 16 genes were affected by ATE in MIA-exposed males, seven of which were modulated in the same direction (all downregulated except for the Gclm gene). Among these, the Ybx1-ps3 pseudogene and Cdkn1a exhibited the highest fold-changes in the PBMC transcriptome. However, since Cdkn1a was also modulated by ATE in both the PBMC and OFC transcriptomes, it is unsuitable as a specific marker of this double-hit vulnerability. Fxyd7 was also among the DEGs modulated in the OFC and PBMC transcriptomes and was not present in the list of genes affected by MIA or ATE in the PBMC transcriptome, warranting further research. See Table S7 for a list of DEGs in PBMCs and OFC and their associated fold changes.

In females, MIA-exposed rats had two genes modulated in both the orbitofrontal and PBMC transcriptomes. One was Gpx3 (upregulated in both cases), coding for glutathione peroxidase 3, and Creg1 (downregulated in PBMCs and upregulated in orbitofrontal tissue), a gene involved in innate immunity and viral response processes. ATE affected only one gene in both the orbitofrontal and PBMC transcriptomes: Nr4a3 (downregulated in PBMCs and upregulated in the OFC), which encodes a member of the steroid-thyroid hormone-retinoid receptor superfamily, potentially acting as a transcriptional activator. Finally, three genes were differentially modulated by MIA depending on ATE experiences: Ciart, Alas2, and Aqp1. Of these, Alas2 and Aqp1 were modulated in the same direction in both the OFC and PBMCs (upregulated in both cases). Alas2 codes for 5’-Aminolevulinate Synthase 2 (an enzyme involved in haem metabolism) and Aqp1 encodes the Aquaporin 1 protein. Both genes had fold-changes of 2 or higher in the PBMCs. Importantly, Alas2 was not modulated in the PBMC transcriptome by either MIA or ATE alone, excluding potential confounding effects of a single-hit experience (Table S7).

## DISCUSSION

Several significant findings are reported in this study. Firstly, even in the absence of overt behavioural disruptions, maternal immune activation (MIA) and adolescent THC exposure (ATE) interacted to affect brain function (as assessed with PET imaging), with these interactions being contingent upon the developmental period examined (early adulthood vs. adulthood) and the sex of the animal. A particularly notable region exhibiting these interactions was the orbitofrontal cortex (OFC). Secondly, subsequent examination of gene expression alterations in the OFC revealed that MIA affected the expression of glutamatergic and serotoninergic receptors, with ATE reversing these effects (i.e., changing from down-regulation to up-regulation) in MIA-exposed rats. Interestingly, ATE alone did not affect these genes. Thirdly, MIA elicited an inflammatory and immune response in the peripheral blood mononuclear cell (PBMC) transcriptome in males, a response that was less pronounced in females. In MIA-exposed rats, ATE altered the gene response direction in some cases, although in others, the differentially expressed genes (DEGs) were also present in the ATE comparison. Finally, we identified several genes with overlapping changes in the brain and periphery that could be explored as biomarkers of vulnerability to schizophrenia in humans.

Additionally, as previously reported, bodyweight loss following an immune challenge (LPS injection) during pregnancy is predictive of the performance of offspring in a series of tests designed to assess symptom dimensions of schizophrenia. When analysing the behavioural performance of the offspring in the selected tests, no clear interactions between MIA and ATE were observed, except for a surprising increase in social novelty preference in animals administered THC during adolescence, notably only in those previously exposed to MIA.

### Lack of behavioural alterations due to MIA, ATE or their combination

No clear behavioural disruptions due to MIA, ATE, or their combination were found at the time of testing. Working memory, as assessed using the Y maze, remained unaltered, although a sex effect was observed, indicating that females generally performed worse on this task. A closer examination of the data suggests that this effect may be driven by slight impairments induced by MIA or ATE, which could be reversed by the combination of both factors; however, these modulations did not reach statistical significance. Previous research demonstrated that LPS-induced MIA was associated with impaired working memory in males in a T maze (Santos-Toscano et al., 2016) but the LPS dose used in that study was higher (100 μg/kg) compared to this study (50 μg/kg). Additionally, animals were tested at slightly different ages (PND60 and PND70, respectively). For a more comprehensive understanding of the effects of early life challenges on working memory, future studies should employ tasks based on operant conditioning, as suggested for emotional regulation deficits (Stupart et al., 2023).

While no alterations in social exploration were observed, an intriguing interaction between MIA and ATE was noted regarding social novelty preference. ATE increased this parameter but only in rats exposed to MIA in utero. Although this result may seem counterintuitive, similar unexpected interactions between risk factors for schizophrenia (such as maternal separation) or genetic modifications of schizophrenia-related genes (such as Comt or Nrg1) have been reported, where the first hit mitigates the deleterious effects of adolescent cannabinoid exposure (Dunn et al., 2020).

There were no significant effects on sucrose preference, prepulse inhibition (PPI), or conditioned taste aversion. Previous studies have shown that LPS-induced MIA could reduce sucrose preference, but these studies were conducted in mice, using a higher dose of LPS, with MIA occurring later in development compared to this study (Rahimi et al., 2020; Ranaei et al., 2020). Consequently, factors such as immunogen dose, species, and timing of the immune challenge appear to be crucial variables influencing the effects of MIA (Boksa, 2010). Regarding PPI, it has been shown that LPS-induced MIA is not associated with PPI deficits (Capellán et al., 2023, 2022) unlike other studies from our laboratory and others that have reported consistent PPI deficits (Santos-Toscano et al., 2020, 2016), depending on the timing of LPS administration, route of administration, PPI protocol, or age at PPI testing. Conditioned taste aversion learning was also assessed due to the lack of previous data on the effects of MIA or adolescent cannabinoid exposure on this form of implicit learning. Additionally, given that some promising antipsychotic drugs, such as TAAR1 agonists, produce conditioned taste aversion (Liu et al., 2022; Shabani et al., 2023), it was deemed crucial to determine whether conditioned taste aversion may be enhanced in this animal model of the disease.

In summary, there were no overt behavioural impairments indicative of manifest symptomatology in this model, at least not at the tested age. To determine whether this absence of disruptions was paralleled by a lack of brain alterations, we proceeded with positron emission tomography (PET) scans.

### MIA and ATE interactions affecting brain activity depending on the sex and age of the animals

ATE alone had only marginal effects on brain metabolism, consistent with previous findings using a lower dose of THC and a different administration regimen (Orihuel et al., 2023) (Figure S4). Interestingly, MIA had a more widespread effect—short-lived in females and with a delayed onset in males—with notable hyperactivations in the frontal cortex, particularly in females (Figure S5). However, the most intriguing findings were the interactions between MIA and ATE. Specifically, we observed a sex- and time-dependent effect of the combination of MIA and ATE on OFC activity. At early adulthood (PND90), female but not male rats with both hits showed aberrant activity, which normalised thirty days later. Other effects were found at PND90, such as decreased activity after exposure to both hits in the thalamus in females and the cortex in males, some of which disappeared by PND120, while others persisted, such as in the pons or cerebellum. Furthermore, additional alterations appeared only later in life (PND120) without detectable presence at earlier periods (PND90), including the amygdala and pons in males. Notably, the most persistent effects in females, sometimes bilateral, were found in the caudal brain regions, such as the pons and cerebellum.

To date, this is the first study to explore the effects of maternal immune activation and adolescent cannabinoid exposure on brain activity using PET. Previous research demonstrated that MIA combined with another adolescent stressor (sub-chronic stress) had significant interactive effects on brain activity assessed by PET, with MIA preventing some long-term alterations in the hippocampus induced by stress (Capellán et al., 2023). However, there was no prior literature on how MIA interacts with adolescent cannabinoid exposure in rats at this level. The effects observed here are unlikely to be attributed to changes in CB1 receptor expression induced by MIA, at least not at the level of the OFC, as MIA does not alter binding levels to these receptors (as measured using a CB1 receptor PET ligand and PET scanning) in the prefrontal cortex, including the orbitofrontal region (Verdurand et al., 2014). These effects were not long-lasting, as the OFC alterations disappeared by PND120. However, other effects emerged at this later age, such as increased activity in the striatum and cortex or hippocampus when both hits were combined, suggesting a dynamic progression of symptoms as the disease advances. Indeed, some findings in humans indicate that orbitofrontal and striatal alterations could differentially predict the emergence of negative symptoms at different stages of the disease spectrum (Kirschner et al., 2021). In males, positive interactions emerged later in life and were confined to the amygdala and pons, with the former having potential implications for affective symptoms that may develop later in life (Potvin et al., 2016). The negative interactions (decreased activity when both hits were combined) were generally restricted to caudal regions (involving the pons and the cerebellum) and were more pronounced in females, especially at PND120. This decreased activation in both regions may be associated with the pons-to-cerebellum hypoconnectivity observed in schizophrenia patients, which correlates with the severity of hallucinations (Abram et al., 2024). In males, there was hypoactivation of the visual cortex in rats exposed to both hits, evident at PND90 but resolved by PND120. This finding in our animal model provides further causal evidence for the visual cortex dysfunctions reported in people with schizophrenia (Schultz et al., 2013).

### Cortical gene expression effect related to schizophrenia

Given the observed sex- and time-dependent effects in the OFC and the noted alterations in this structure in schizophrenia (Chen et al., 2022; Nakamura et al., 2019, 2007), we delved deeper into the neurobiological mechanisms involved, performing RNA sequencing on orbitofrontal samples from rats exposed to MIA and ATE.

MIA downregulated several genes related to glutamatergic and serotoninergic receptors implicated in schizophrenia, such as Gria3 and Grin2a (Singh et al., 2022; Trubetskoy et al., 2022) and Htr2a (García-Bea et al., 2019; Kang et al., 2020) among others. Notably, ATE in MIA rats shifted this gene expression modulation from down-regulation to up-regulation. ATE alone did not affect these genes, suggesting that the effects observed when both hits occur in the same animals are not merely due to an exacerbation of adolescent cannabinoid exposure effects. Grin2a has strong evidence of involvement in schizophrenia (Singh et al., 2022), and its up-regulation may indicate a compensatory response to potential receptor hypofunction, consistent with the NMDA receptor hypofunctionality hypothesis (Nakazawa and Sapkota, 2020). Similarly, the Errb4 gene, which codes for a receptor tyrosine kinase involved in NMDA receptor function regulation (Hahn et al., 2006; Vullhorst et al., 2015), was up-regulated in two-hit rats and down-regulated by MIA. Interestingly, ATE alone did not affect these glutamatergic or serotonergic genes, either in males or females. ATE resulted in a notable sex-dependent effect, with the neural cell adhesion molecule 1 gene (Ncam1) being up-regulated in females and down-regulated in males. Neural cell adhesion molecules are vital for central nervous system development (Sytnyk et al., 2017), and the Ncam1 gene has been identified as a hub for the shared genetic basis of cannabis, cigarette smoking, and schizophrenia (Song et al., 2022). This suggests that ATE may confer a sex-specific vulnerability to schizophrenia through specific modifications of central nervous system developmental programmes.

### Gene expression changes in PBMCs that parallel the effects found in the brain

In an effort to identify potential markers of schizophrenia susceptibility, we conducted RNA sequencing on PBMCs obtained from male and female rats exposed to MIA, ATE, or their combination. PBMCs have been suggested as potential candidates for biomarker discovery in schizophrenia (Astafeva et al., 2023) including for identifying first-onset cases as opposed to differentiated schizophrenia patients (Zaki et al., 2022). We focused on genes modulated in both the OFC and PBMCs, differentially affected by ATE depending on MIA experience. Among these, Cdkn1a emerged as a gene with high potential for exploration as a biomarker, modulated in the same direction in both the OFC and PBMC transcriptomes and among those with higher fold-change values. The expression of this gene is modulated in schizophrenia (Merikangas et al., 2022), making it a promising biomarker candidate. However, this gene is also modulated by ATE in PBMCs (and in the OFC in an opposite direction), complicating its potential specificity and sensitivity as a biomarker, especially considering a history of cannabis use. Another candidate is the Fxd7 gene, modulated in the same direction in both PBMCs and OFC and only in the OFC by ATE, allowing it to be measured in PBMCs without potential confounds. Fxd7 is a Na-K ATPase that regulates neuronal excitability in the brain (Yap et al., 2021), and the unique pattern of differences in its expression suggests it could be a reliable marker in PBMCs.

In females, the Alas2 gene was modulated in the same direction in both PBMC and OFC transcriptomes by ATE, depending on MIA experience, and was not modulated in PBMCs by either hit alone. Moreover, it showed a high fold change (2.49) in PBMCs, making it a promising peripheral biomarker candidate. The Alas2 locus, located in the Xp11 chromosomal region, was identified as a marker for schizophrenia in a linkage study (Dann et al., 1997).

## Conclusions

We have provided comprehensive evidence for the causal role of the combined effects of MIA and ATE in creating a brain susceptibility status compatible with an increased risk of developing schizophrenia, even in the absence of overt behavioural manifestations of the disease. The lack of behavioural alterations despite severe brain modifications highlights the need for more sensitive behavioural assays for modelling symptom dimensions of the disease. Importantly, the OFC appears to be a critical region in these effects. Additionally, this study further supports the role of glutamatergic transmission in schizophrenia (with a significant role for the Grin2a gene) and suggests specific targets for sex-specific biomarker discovery studies. Furthermore, we confirm previous evidence that weight loss associated with prenatal infections could be an important predictor of the behaviour of the progeny, excluding the contribution of other potential indices.

In conclusion, we suggest that cannabis consumption during adolescence may indeed trigger schizophrenia but only in individuals with prior vulnerability, such as maternal immune activation during gestation.

## Acknowledgements

This study was supported by grant PID2019-104523RB-I00 to AHM funded by MICIU/AEI/ 10.13039/501100011033, and grants from the Ministry of Health-Plan Nacional sobre drogas (National Plan on Drugs): 2021I039 and EXP2022/008739 (supported by the Plan for Recovery, Regeneration and Resilience funded by the European Union - NextGenerationEU).

We would like to thank Marta Casquero-Veiga for the support with SPM analyses.

