## Supplementary materials and methods for "A hidden mark of a troubled past: neuroimaging and transcriptomic analyses reveal interactive effects of maternal immune activation and adolescent THC exposure in the absence of overt behavioural disruptions"

**SUPPLEMENTARY INFORMATION**

**SUPPLEMENTARY MATERIALS AND METHODS**

**Animals**

All experiments were approved by the Ethics Committee of UNED and the Autonomous Community of Madrid (PROEX 177.4/20). The animals were always maintained and treated in accordance with the European Union Laboratory Animal Care Guidelines (EU Directive 2010/63/EU), and the "Principles of Animal Welfare" were meticulously followed.

All the animals were Sprague-Dawley (SD) CD rats obtained from Charles-River. The animals were always kept at a constant temperature (23ºC ± 1ºC) and humidity (50% ± 10) and on an inverted 12-hour light cycle (lights on at 8:00 PM). Unless otherwise specified, the animals always had ad libitum access to food (commercial rodent diet Rod14; Sodispan Research; Madrid), tap water, and environmental enrichment. The behavioural experiments described were always conducted during the animals' active cycle, between 9:00 AM and 7:00 PM, with lighting between 20 and 50 lux.

**Mating and LPS administration**

Females were mated with males at a 1:1 ratio. Daily vaginal smears were taken to analyse the presence of sperm and establish gestational day (GD) 0, at which point the female was separated from the male. From this point until the day of delivery, weights were recorded daily to monitor the progress of gestation.

On GD15 and GD16, the rats were injected intraperitoneally, each day, with a dose of 50 µg/Kg of lipopolysaccharides (LPS) 0111:B4 (Sigma-Aldrich) dissolved in sterile saline solution (0.9% NaCl solution; Braun) (n=11) or with saline (n=12), both at a volume of 1 mL/Kg. Rectal temperatures were recorded with rectal probes (Bioserve) immediately before the injections and 60, 120, and 240 minutes afterward as an indicator of MIA. Bodyweight was measured on both injection days and on two additional consecutive days (from GD15 to GD18).

At birth, pups were marked with an ink tattoo on their hind paws to identify group allocation. All litters were redistributed among LPS- and saline-exposed dams to ensure that each raised a similar number of pups from each group. For the redistribution of pups, litters with more than 1 day of difference between their birth dates were not mixed. Additionally, it was ensured that each litter had no more than 2 pups' difference between males and females.

All pups were weaned on postnatal day (PD) 22 and grouped according to sex and prenatal exposure. Subsequently, they were assigned to THC and vehicle groups, ensuring that each group included pups from all mothers.

**Adolescent THC exposure**

Δ^9^-Tetrahydrocannabinol (THC) was obtained from THCPharm (Frankfurt, Germany). The resin was dissolved in pure ethanol (30 mg/mL) and stored as a stock solution at -30ºC. Working solutions were prepared daily by mixing the stock solution with castor oil PEG-35 (Kolliphor; Merck) and saline solution in a ratio of 1:1:18. All solutions were prepared in siliconized vials (Sigmacote; Sigma) under a nitrogen-saturated atmosphere and protected from light. Vehicle group aliquots were prepared in the same manner but without adding THC to the stock solution. In all cases, the ethanol administered together with the THC was below 0.1 g/Kg, a dose that does not have significant effects on behaviour (Frye & Breese, 1981).

Increasing doses of THC were administered via intraperitoneal route at a volume of 2 mL/Kg. Between PD30 and PD32, a dose of 2.5 mg/Kg was administered; between PD33 and PD36, the dose was increased to 5 mg/Kg, and finally, between PD37 and PD40, the dose was 10 mg/Kg (protocol modified from Rubino et al., 2008).These doses mimic heavy marijuana use, as well as the increasing pattern of consumption seen in human adolescents (Zamberletti et al., 2014).

**Early markers in the dams of behavioural deficits in the offspring**

In order to find early markers in the female progenitors that may help predict alterations in offspring, prior to mating we performed an assessment of behaviours related to anxiety and reactivity in new environments. Additionally, the physiological and immune reactivity to LPS of the dams during gestation and after weaning was explored.

Elevated plus maze

Anxiety-like behaviour was assessed approximately one week before mating, at an age of 10-12 weeks. Firstly, we used the elevated plus maze which is based on rodents' innate tendency to explore their environment, avoiding open, bright, and elevated places (Campos et al., 2013; Handley & Mithani, 1984). For this purpose, we used a cross-shaped maze made of black methacrylate and expanded PVC (Forex®), elevated 50 cm from the floor, with 2 arms with high walls (closed arms (50x10x50cm)), and two arms without walls, (open arms (50x10cm). The test duration was 5 minutes and was carried out as described in (Higuera-Matas et al., 2009). The data were analysed using a formula similar to the protocol validated by (Pellow & File, 1985):

$${Index}_{EPM}=\frac{\left( Time in open arms \right)-(Time in open arms)}{(Time in open arms) + (Time in closed arms)}$$

Locomotor activity

Two days after the EPM, we analysed locomotor activity in a novel environment using an open field maze (50x50x50cm) made of methacrylate (Himanshu et al., 2020). Rats were placed in the centre of the maze and allowed to explore for 40 minutes. We used total distance travelled as a measure of locomotor activity.

Immune reactivity and cytometry assay

The magnitude of the immune response to LPS was analysed in all dams approximately two weeks after weaning. For this purpose, on the first day, half of each group was injected with LPS (50 µg/Kg, i.p.) and the other half with saline solution. After a day of rest, the procedure was repeated by injecting LPS or saline to those who had not received it before. Rectal temperature (taken just before injection, after 60min, 120min, and 240min) and the distribution of PBMC populations were used as measures of this immune response.

Monocyte, B lymphocyte, and T lymphocyte populations of PBMC were analysed by flow cytometry. A volume containing 1x10^6^ cells was incubated with primary antibodies (Table S1) diluted in PBS and 10% fetal bovine serum for 1 hour at room temperature. After 3 washes with PBS, the cells were fixed with FACS™ Lysing Solution (BD Biosciences) and analysed on a FACSCalibur flow cytometer (Becton Dickinson) using conventional techniques.

| **Antibody** | **Fluorophore** | **Population** | **Concentration** | **Manufacturer** | **Reference** |
| --- | --- | --- | --- | --- | --- |
| **CD3** | FITC | T lymphocytes | 2.5 µL/mL | Becton Dickinson | 557015 |
| **CD11b/c** | PE | Monocytes | 2.5 µL/mL | Becton Dickinson | 554862 |
| **CD45RA** | PECy5.5 | B lymphocytes | 2.5 µL/mL | Becton Dickinson | 557015 |

**Table S1:** **Main features of the antibodies used to detect the main immune cell populations in the dams after saline or LPS injection.**

**Behavioural Tests in Offspring**

When the animals reached adulthood, a comprehensive set of behavioural tests was initiated to evaluate behaviours that could be similar to observed in the different symptoms of schizophrenia. To assess potential cognitive alterations, we evaluated working memory (Y maze) and pre-attentional filtering (prepulse inhibition of the acoustic startle response). For negative symptoms, we analysed social interactions and sucrose preference as measures of anhedonia. Finally, we evaluated conditioned taste aversion as a form of implicit learning which could also modify the feasibility of some new antipsychotic medications. Initial group sizes are as follows: (males: saline+vehicle:n=13; LPS+vehicle: n=11; saline+THC: n=11; LPS+THC: n=11; females: saline+vehicle: n=11; LPS+vehicle:n=10; saline+THC: n=9; LPS+THC: n=9)

Working Memory

The Y-maze has traditionally been used to evaluate working memory (Conrad et al., 1997; Lainiola et al., 2014; Prieur & Jadavji, 2019). Here, we used the continuous variant of the Y-maze test, which consists of allowing the animals to freely explore in order to record the sequence of arm entries. This test is based on the tendency of rodents and other animals to alternate arm exploration in the absence of reinforcers and, therefore, on the need to remember the recently explored arms (Hughes, 2004). The Y-shaped maze was made of methacrylate with a long arm (22x42.5x14cm) and two short arms (22x38.5x14cm). Approximately on PND71, the animals were placed at the end of the long arm facing away from the maze centre and were allowed to explore the maze for 10 minutes (Miedel et al., 2017) . Both the entries into each arm and the sequence of entries were recorded. We used the percentage of alternation, calculated by dividing the number of triads (3 consecutive entries) into different arms by the maximum possible number of triads (total entries minus 2) as a measure of working memory.

Social Interaction

In this work, we have focused on two components of social interaction: sociability and social novelty preference. We used a conventional three-chamber arena made of black methacrylate (Yang et al., 2011) with a central chamber (39.5x79x29.5cm) connected to two larger lateral chambers (39.5x79x39cm) by two guillotine doors.

First, the animals were allowed to habituate to the central chamber for 3 minutes with the guillotine doors closed. After the adaptation period, two cylindrical containers (30cm high and 13.5cm in diameter) were placed in the centre of each lateral chamber. The containers had interspaced bars as walls, which allowed the contact between individuals. In phase 1, one of the cylinders contained a conspecific rat of the same sex (Conspecific 1) while in the other container we placed an inanimate object (a stuffed rat). Phase 1 was conducted immediately after habituation; the guillotine doors were opened, and the rat was allowed to explore the entire maze for 8 minutes.

At the end of phase 1, the rat was relocated to the central chamber, and a new conspecific rat of the same sex (Conspecific 2) was placed in the cylinder that contained the inanimate object. The doors were open again, and the behaviour of the test rat was recorded for another 8 minutes. In both cases, interaction times were analysed, defined as the time the animal's head was within a 5 cm radius of the cylinder and facing it. Sociability and social novelty indices were calculated as follows:

Sociability: $\frac{Interaction with Conspecific1-Interaction with inanimate object}{Total Time}$

Social novelty:$\frac{Interaction with Conspecific2-Interaction with Conspecific 1}{Total Time}$

Test for incidental associations and conditioned taste aversion

We used 50mL Falcon™ tubes as drinking bottles, which were weighed before and after the tests. Rubber stoppers fitted with a metal tube were also used to allow the animals to drink.

Throughout this procedure the animals had 1 hour of water access in the morning, with the scents and/or flavors dissolved or not depending on the experimental phase.

We first allowed a drinking training phase, so that the animals learned to drink from the new drinking bottles. The criterion was that all animals drank at least 3mL in one session or a minimum of 3 days. Then, the animals were exposed to 6 sessions of compound exposure (Sensory Preconditioning Phase). Each session consisted of two days, during which a 1-hour exposure was carried out in the morning to one of the two possible odor-taste combinations: 0.01% almond odor (benzaldehyde; Sigma) with 5 mM NaCl taste (VWR) and 0.05% banana odor (isoamyl acetate; Sigma) with 0.316 mM HCl (PanReac Applichem). The odors were presented in the water itself as flavourings, and their concentrations were 0.05% almond (benzaldehyde) and 0.017% banana (isoamyl acetate). The concentrations of odors and flavors were chosen so that there were no preferences (Busquets-Garcia et al., 2017; Tordoff et al., 2008) and were adjusted accordingly. The order of presentation of both combinations was counterbalanced among animals.

After the sensory preconditioning phase, the conditioning phase took place. In the morning, all experimental groups were exposed 1 hour to one of the odours (the one they had on the first day of sensory preconditioning), and 5 minutes later, they were administered an injection of LiCl (5 mL/Kg; i.p., 0.3 M in 0.9% NaCl; Sigma) considered aversive (Gore-Langton et al., 2015). The next day, the procedure was repeated with the other odour using a saline injection as control. There were 3 conditioning sessions. After this phase, the animals had a recovery day with 1 hour of water access in the morning.

After the recovery phase, the test phase began, which in all cases consisted first of the mediated test and then the direct test. There was one test day was in which both tastes were presented through a preference test (prior exposure to both options) for 1 hour. The mL consumed in both cases were recorded.

In the mediated test we tested whether there was an aversion to the flavour that had been preconditioned with the odour paired with LiCl. In the direct test, we checked whether direct aversive conditioning had occurred. A preference test (prior exposure to both options) between the two odours dissolved in water was conducted. The mL consumed from each bottle were recorded. Animals were single-housed for this test. Both odours, despite being in closed bottles, were always presented in opposite parts of the room. Due to an error in the allocation of the bottles, a side-bias developed in the mediated test and therefore we will only report results of the direct conditioning test. However, the study of incidental associations in this paradigm should be studied further (see (Moreno-Fernández et al., 2024)

Sensory-motor gating

The prepulse inhibition of the startle response was used as a measure of sensory-motor gating. We employed a methacrylate box (28x15x17cm) with a vibration-sensitive platform inside (Cibertec), calibrated daily using a 200g weight. The test lasted approximately 26 minutes and consisted of a previously validated sequence of trials (Capellán et al., 2022). Prepulses of 4 or 12 decibels (dB) above background noise (65 dB) with intervals of 30 or 120 milliseconds (ms) between prepulse and pulse (120 dB) were used. Prepulse inhibition was calculated using the following formula, considering the peak startle amplitude:

%PPI=$100-((\frac{mean of prepulse trials}{mean of pulse trials})*100)$

Habituation was calculated by exposing the animals to 6 trials with only the main pulse at the beginning and end of the session (not included in the PPI calculations). The percentage of habituation was calculated by analysing the startle amplitudes as follows:

%Habituation=$(\frac{Mean startle first block of trials-Mean last block of trials}{Mean first block of trials})*100$

Anhedonia

To analyse the potential existence of anhedonia, the sucrose preference test was used based on the literature (He et al., 2020). We restricted access to water to 1 hour in the afternoon in their home cage, and the experiments took place in the morning. The rats had 4 daily sessions of 1 hour where they were individualized in cages with access to two water bottles (one on each side) with tap water. After this habituation phase, 4 new sessions were conducted, replacing the water in one of the two bottles with 1% sucrose (Sigma) prepared on the same day. The position of the bottles was counterbalanced among animals and days. To measure potential anhedonia, the percentage of sucrose preference was used:

%Preference=$\frac{Volume of sucrose consumed}{volume of sucrose+volumen of water consumed}*100$

**Positron Emission Tomography**

The effects of MIA ATE, and their interaction on [18F]-2-fluoro-2-deoxy-D-glucose (18F-FDG) brain uptake were studied in 80 Sprague Dawley rats (40 males and 40 females) divided in 4 groups (n= 10) at two different time points: postnatal day 60 (PET1) and 120 (PET2)

Positron emission tomography/computed tomography (PET-CT) studies were performed using a small-animal SuperArgus PET/CT (SEDECAL, Madrid, Spain). 16 h fasted animals and anaesthetised with 2–3% isoflurane in 100% medical oxygen (1 L/min) were scanned for 30 min after administration of 14.06 +/- 3.58 MBq of 18F-FDG via the tail vein. During the PET studies (energy window 350–700 KeV and 45 min list mode acquisition) and CT (voltage 30 kV, current 500 μA, 8 shots, 360 projections and standard resolution) its temperature was maintained at 37ºC. 3D-OSEM (ordered subset expectation maximization) algorithm (21 subsets and 5 iterations) imagen reconstruction was accomplished, corrections for random and scatter were applied.

PET image post-processing was performed using a previously published protocol (Soto-Montenegro et al., 2014) adapted to PMOD software (PMOD Technologies GmbH, Version 4.302). Briefly, the reconstructed images were spatially registered using elastic transformations with an automatic algorithm based on mutual information. PET intensity values were normalised to the average uptake in the whole brain and then, images were smoothed with a Gaussian kernel 2 times the voxel size at full width at a half maximum (FWHM). Automatic registration between PET and MRI scans was not feasible. Therefore, a manual registration process employing PMOD software was used. Then, a brain mask obtained from the MRI was applied over each PET scan to exclude extra-cerebral areas.

Statistical analysis was performed using SPM software (http://www.fil.ion.ucl.ac.uk/ spm/software/spm12/). A two-way ANOVA (p.value < 0.01 uncorrected) was conducted to examine the effects of THC and LPS on 18F-FDG consumption in the brain for each sex (male and female) and time point (PET1 and PET2). Only clusters with p.value <0.05 uncorrected were considered to minimise the effect of type I errors.

**RNA Extraction**

At around PD90, rats were lightly anesthetised with isoflurane and killed by decapitation. The brains were then carefully extracted and frozen at -80ºC. For dissection the brains were thawed but kept ice cold. Slices were obtained using acrylic matrices and razor blades. Using the Paxinos & Watson, 2007 atlas as a reference, the orbitofrontal cortex (OFC) was dissected from the corresponding slices, weighed, and kept on dry ice until storage at -80ºC. All surfaces and dissection equipment were treated with RNaseZap (Thermo) to digest RNases.

For RNA extraction, the samples were homogenized in homogenization buffer (50mM HEPES, pH 7.5, 320mM sucrose, 20mM sodium butyrate, protein and phosphatase inhibitors in DEPC water). RNA extraction was performed using a column extraction with the RNeasy Mini Kit (Qiagen) and RNase-Free DNase Set (Qiagen) for on-column DNA digestion. The concentration and RIN (RNA integrity number) were obtained using an Agilent Bioanalyzer 2100 with the RNA 6000 nano kits (Agilent). We selected 3 rats of each group for sequencing based on RIN and RNA concentration.

**Massive RNA Sequencing**

Massive RNA sequencing was performed by the Genomics Unit of the Madrid Science Park. Libraries were prepared using the NEBNext® Poly(A) mRNA Magnetic Isolation Module and NEBNext® UltraTM II Directional RNA Library Prep for Illumina® kits (New England Biolabs), starting from 91 ng of total RNA. The instructions from "Chapter 1: Protocol for use with NEBNext® Poly(A) mRNA Magnetic Isolation Module" were followed. For library amplification as mentioned in the protocol, a 13-cycle PCR was used. For PBMCs, libraries and samples were validated and quantified using the Agilent Bioanalyzer 2100 and Agilent RNA 6000 and Agilent High Sensitivity kits. In the case of the orbitofrontal cortex, both samples and libraries were validated and quantified using the 4200 TapeStation System (Agilent) and RNA Screen Tapes and D1000 Screen Tapes kits. After library quantification, an equimolar mixture was prepared, which was purified with AMpure XP beads (Beckman Coulter) and quantified by qPCR using a reference library from the Genomics Unit and the KAPA SYBR® FAST qPCR kit for the LightCycler® 480 instrument. The equimolar mixture was sequenced on the Illumina® NovaSeqTM 6000 instrument using the NovaSeqTM Reagent 1.5 kit (100 cycles) in a single-read 1x100 run.

After sequencing, the TopHat program included in the G-PRO suite (Biotechvana) was used to map and locate the different sequences against the reference genome (Rattus norvegicus Rn6.0.96), obtaining a percentage of correctly mapped reads ranging from 93 to 95%. Fragments of 19,186 and 21,437 genes were found in PBMC and OFC sequencing, respectively, and a global mean fragment cutoff of less than 240 and 360 was applied as an exclusion criterion. This left a total of 11,855 and 13,942 genes considered expressed in each region. Subsequently, a differential expression analysis was performed using the CuffDiff program, which analyzes the RNA expression of each gene, normalized by its size and the global RNA expression of each sample, and performs a comparison between groups applying false discovery rate (FDR) correction and obtaining Fold Change (FC) values.

Sequencing data have been deposited in the GEO database under the accession number GSE273695.

**Data Analysis**

Video Analysis

Animal behaviour was analysed using the ANY-maze Video Tracking System v.6.32 (Stoelting Co.) or higher.

Indices and Correlations

As early markers that could predict some of the deficits caused by one or both impacts, a predictive index for mothers (IM) was calculated using normalized data from the elevated plus maze, locomotor activity, the greatest rectal temperature variation after LPS administration, weight loss after the first LPS injection and the percentage change in T lymphocytes after an LPS injection. This index was calculated as follows:

𝐼𝑀 = −Normalized LCE − locomotor activity − temperature variation + weight loss − %Δ T lymphocytes

Similarly, an index was calculated to group the results of all offspring behavioural tests (IC). For this purpose, the mean of each litter normalized Z results for each test was used:

𝐼𝐶 = −Memory − Sociability − Novelty − PPI − Preference scores

Once both indices were obtained, several correlation analyses were performed between both indices and their components.

Data Normalization

To minimize variability between tests conducted in several batches of animals across time, we normalised the data using Z-scores. For Z transformation, the mean and standard deviation of all animals in the corresponding batch were used for each transformed variable. For a more understandable visualization of the data, they were transformed back again to direct scores before graphical representation. For this purpose, each Z-scored data point was multiplied by the joint standard deviation of the raw data from the included batches and the mean of these batches was subsequently added.

To ensure comparability of all index components, they were normalized with Z. The mean and standard deviation of all animals in the batch to which they belonged were used for each variable. In offspring, after normalization, the mean of each litter was obtained to be compared with the mother's index. Since each litter contained males and females, both THC and vehicle, whenever possible, the mean of each group within each litter was also calculated to correlate with its mother, to find presence or absence of correlations (e.g., THC-males correlations with their mothers or vehicle-females correlations with their mothers).

The fragments from the massive RNA sequencing were normalized using the DESeq2 package of RStudio v.2022.12.0+353 (Posit Software, PBC) and R Statistical Software v.4.2.2 (R Core Team 2022). This method considers the total number of fragments from each sample and allows comparisons between them. Finally, the normalized fragments were transformed into Z-values using the mean and standard deviation of the normalized fragments for each gene. With these values, heatmaps were generated to allow comparison of each gene separately between the different samples.

Statistics

IBM SPSS Statistics v.26.0 was used for statistical analysis of the data. All results are expressed as mean ± standard error of the mean (SEM) unless otherwise stated. Outliers were identified using SPSS's interquartile range criterion with a p-value <0.05. Normal distribution and homoscedasticity of the data were evaluated using the Kolmogorov-Smirnov and Shapiro-Wilk tests for normality and the Levene test for equality of variances. Square roots, natural logarithms, or inverse transformations were used to correct distributions or lack of homoscedasticity of the data when necessary. Unless otherwise stated, experiments were analysed using independent 3-way ANOVA considering "Sex" (male or female), "MIA" (saline/LPS), and "ATE" (vehicle or THC) as between-subject factors. Mixed 3-way ANOVA was used to analyse weights during THC exposure and 18F-FDG uptake using exposure days ("Days"). They were also used to analyse baseline preferences in the three-room maze (inanimate object vs. conspecific1 and conspecific1vs. conspecific2), sucrose preference (water vs. sucrose), and preferences for dCS (+ vs. -). Greenhouse-Geisser corrected F values were used, and significant interactions were analysed by simple effects analysis. F values, effect size (partial eta squared η2p), were indicated whenever necessary.

**SUPPLEMENTARY RESULTS**

|  | **Effect (ANOVA)** | ***F*(dfM, dfR)** | ***p-value*** | **η^2^p** |
| --- | --- | --- | --- | --- |
| Temperature GD15 | **Time*MIA** | *F*(2, 27) =16.28 | 0.000 | 0.538 |
|  | **Time*[MIA=LPS] (T0 vs T60)** | *F*(1, 7) =6.01 | 0.044 | 0.462 |
|  | **Time*[MIA=LPS] (T0 vs T120)** | *F*(1, 7) =49.67 | 0.000 | 0.876 |
|  | **Time*[MIA=LPS] (T0 vs T240)** | *F*(1, 7) =12.01 | 0.010 | 0.632 |
| Temperature GD16 | **Time** | *F*(1, 19) =3.98 | 0.047 | 0.235 |
|  | **Time (T0 vs T60)** | *F*(1, 13) =0.93 | 0.351 | - |
|  | **Time (T0 vs T120)** | *F*(1, 13) =8.63 | 0.012 | 0.005 |
|  | **Time (T0 vs T240)** | *F*(1, 13) =5.70 | 0.033 | 0.055 |
| Bodyweight GD15-GD18 | **GD** | *F*(2, 27) =110.01 | 0.000 | 0.887 |
|  | **GD*MIA** | *F*(2, 27) =11.13 | 0.000 | 0.443 |
|  | **GD*[MIA=LPS]** | *F*(1, 8) =38.22 | 0.000 | 0.845 |
|  | **GD*[MIA=LPS] (GD15 vs GD16)** | *F*(1, 7) =4.67 | 0.067 | - |
|  | **GD*[MIA=LPS] (GD15 vs GD17)** | *F*(1, 7) =0.17 | 0.689 | - |
|  | **GD*[MIA=LPS] (GD15 vs GD18)** | *F*(1, 7) =29.66 | 0.001 | 0.809 |
|  | **GD*[AIM=sal]** | *F*(2, 12) =109.04 | 0.000 | 0.94 |
|  | **GD*[AIM=sal] (GD15 vs GD16)** | *F*(1, 7) =51.44 | 0.000 | 0.880 |
|  | **GD*[AIM=sal] (GD15 vs GD17)** | *F*(1, 7) =50.21 | 0.000 | 0.878 |
|  | **GD*[AIM=sal] (GD15 vs GD18)** | *F*(1, 7) =198.45 | 0.000 | 0.966 |

**Table S2:** **LPS effects in the dams:** statistics for the different effects caused by LPS injection on the core body temperature and body weight of the dams. Interactions are expressed with “*” between the corresponding factors. In simple effects, the analysed group appears in brackets and the contrasts in parentheses. Between-subject effects are presented with the name of the factor alone (e.g.: MIA). Within-subject factors: Time (minutes) and GD (day). Between-subject factors: MIA (saline/LPS).

|  | **Effect (ANOVA)** | ***F*(dfM, dfR)** | ***p-value*** | **η^2^p** |
| --- | --- | --- | --- | --- |
| Y maze | **Sex** | *F*(1, 77) = 7.66 | **0.007** | 0.090 |
|  | **Sex*MIA** | *F*(1, 77) = 0.00 | 0.952 | - |
|  | **MIA** | *F*(1, 77) = 0.00 | 0.947 | - |
|  | **ATE** | *F*(1, 77) = 0.47 | 0.497 | - |

**Table S3: F, p and η2p values for each effect studied in the ANOVA of the Y maze performance.**

|  | **Effect (ANOVA)** | ***F*(dfM, dfR)** | ***p-value*** | **η^2^p** |
| --- | --- | --- | --- | --- |
| Social Interaction | **Sex** | *F*(1, 78) = 1.01 | 0.318 | - |
|  | **MIA** | *F*(1, 78) = 0.13 | 0.719 | - |
|  | **ATE** | *F*(1, 78) = 0.58 | 0.449 | - |
|  | **MIA*ATE** | *F*(1, 78) = 1.18 | 0.281 | - |
| Social Novelty Preference | **Sex** | *F*(1, 78) = 1.05 | 0.310 | - |
|  | **MIA** | *F*(1, 78) = 0.00 | 0.960 | - |
|  | **ATE** | *F*(1, 78) = 0.54 | 0.463 | - |
|  | **MIA*ATE** | *F*(1, 78) = 5.56 | **0.021** | 0.066 |
|  | **[MIA = sal]*ATE** | *F*(1, 83) = 1.32 | 0.254 | - |
|  | **[MIA = LPS]*ATE** | *F*(1, 83) = 5.04 | **0.027** | 0.057 |

**Table S4: F, p and η2p values for each effect studied in the ANOVA of the three-chamber test.**

| **PPI condition** | **Effect (ANOVA)** | ***F*(dfM, dfR)** | ***p-value*** | **η^2^p** |
| --- | --- | --- | --- | --- |
| 12dB_120ms | **Sex** | *F*(1, 78) = 0.29 | 0.591 | - |
|  | **MIA** | *F*(1, 78) = 0.40 | 0.531 | - |
|  | **ATE** | *F*(1, 78) = 0.02 | 0.900 | - |
|  | **Sex*MIA** | *F*(1, 78) = 0.03 | 0.855 | - |
|  | **Sex*ATE** | *F*(1, 78) = 0.13 | 0.719 | - |
|  | **MIA*ATE** | *F*(1, 78) = 0.00 | 0.963 | - |
|  | **Sex*MIA*ATE** | *F*(1, 78) = 0.00 | 0.987 | - |
| 4dB_120ms | **Sex** | *F*(1, 44) = 2.63 | 0.112 | - |
|  | **MIA** | *F*(1, 44) = 0.00 | 0.977 | - |
|  | **ATE** | *F*(1, 44) = 1.14 | 0.291 | - |
|  | **Sex*MIA** | *F*(1, 44) = 1.88 | 0.178 | - |
|  | **Sex*ATE** | *F*(1, 44) = 0.24 | 0.624 | - |
|  | **MIA*ATE** | *F*(1, 44) = 1.65 | 0.206 | - |
|  | **Sex*MIA*ATE** | *F*(1, 44) = 0.43 | 0.514 | - |
| 12dB_30ms | **Sex** | *F*(1, 62) = 0.05 | 0.824 | - |
|  | **MIA** | *F*(1, 62) = 2.70 | 0.106 | - |
|  | **ATE** | *F*(1, 62) = 0.00 | 0.997 | - |
|  | **Sex*MIA** | *F*(1, 62) = 2.58 | 0.114 | - |
|  | **Sex*ATE** | *F*(1, 62) = 0.02 | 0.893 | - |
|  | **MIA*ATE** | *F*(1, 62) = 0.234 | 0.630 | - |
|  | **Sex*MIA*ATE** | *F*(1, 62) = 0.00 | 0.972 | - |
| 4dB_30ms | **Sex** | *F*(1, 47) = 4.59 | **0.037** | 0.089 |
|  | **MIA** | *F*(1, 47) = 0.03 | 0.858 | - |
|  | **ATE** | *F*(1, 47) = 0.26 | 0.613 | - |
|  | **Sex*MIA** | *F*(1, 47) = 0.13 | 0.718 | - |
|  | **Sex*ATE** | *F*(1, 47) = 0.02 | 0.885 | - |
|  | **MIA*ATE** | *F*(1, 47) = 3.05 | 0.087 | - |
|  | **Sex*MIA*ATE** | *F*(1, 47) = 0.014 | 0.905 | - |
| Habituation | **Sex** | *F*(1, 79) = 1.28 | 0.261 | - |
|  | **MIA** | *F*(1, 79) = 1.70 | 0.196 | - |
|  | **ATE** | *F*(1, 79) = 0.11 | 0.747 | - |
|  | **Sex*MIA** | *F*(1, 79) = 3.78 | **0.055** | 0.046 |
|  | **[Sex = M]*MIA** | *F*(1, 83) = 5.82 | **0.018** | 0.066 |
|  | **[Sex = F]*MIA** | *F*(1, 83) = 0.19 | 0.660 | - |
|  | **Sex*ATE** | *F*(1, 79) = 0.41 | 0.523 | - |
|  | **MIA*ATE** | *F*(1, 79) = 0.35 | 0.556 | - |
|  | **Sex*MIA*ATE** | *F*(1, 79) = 0.06 | 0.814 | - |

**Table S5: F, p and η2p values for each effect studied in the ANOVA of the prepulse inhibition test.**

| **Offspring** |  | DI | EPM | Loc. Act. | Temp. | BW | T Lymp. |
| --- | --- | --- | --- | --- | --- | --- | --- |
| OI | Pearson | 0.589 | -0.220 | -0.172 | -0.273 | 0.825 | 0.333 |
|  | *p-value* | 0.125 | 0.601 | 0.684 | 0.513 | **0.012** | 0.420 |
|  | N | 8 | 8 | 8 | 8 | 8 | 8 |

**Table S6: correlation analysis of the offspring index, dam index and its components.**

**Table S7: significant differentially expressed genes in PBMCs and OFC with the gene names and fold change.** See associated excel file containing table S7.

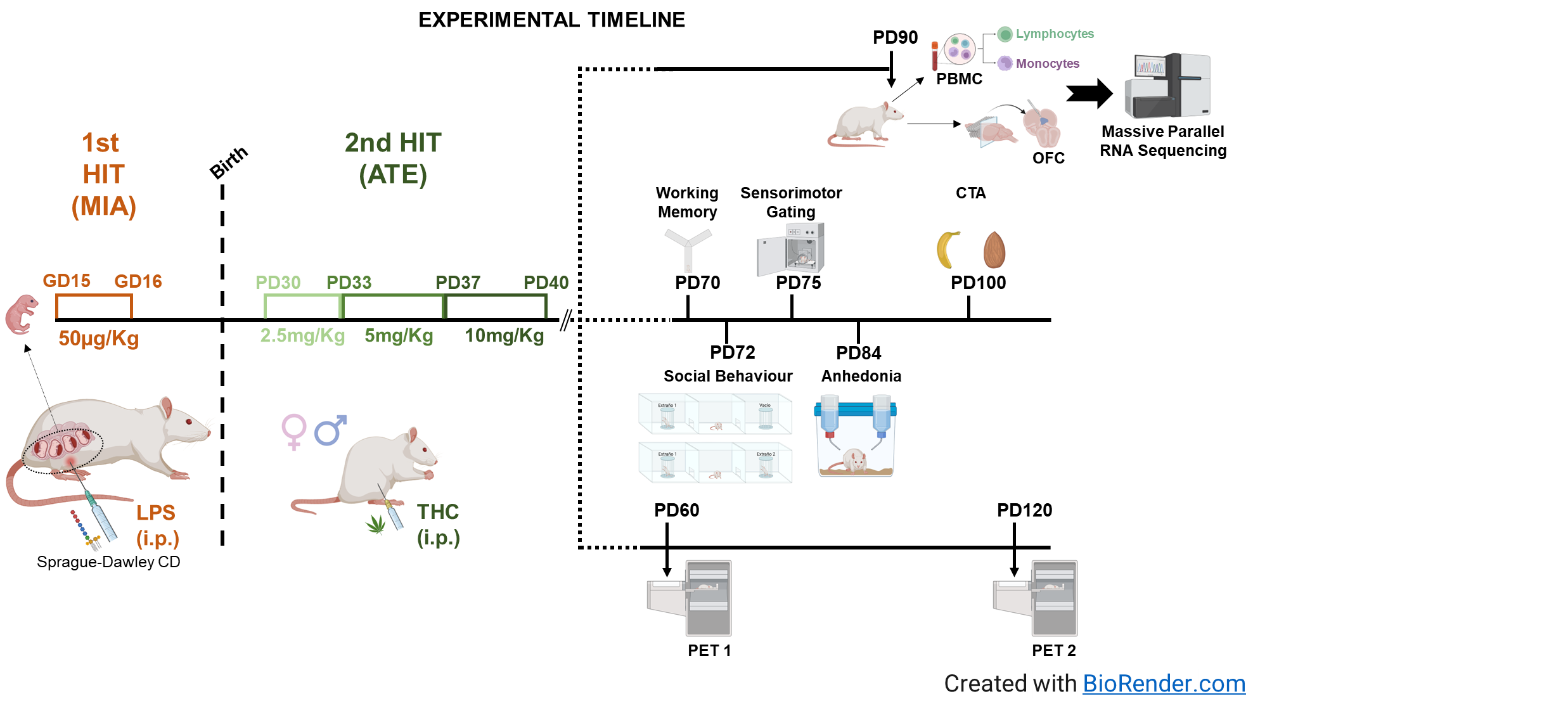

**Figure S1: Experimental Timeline**

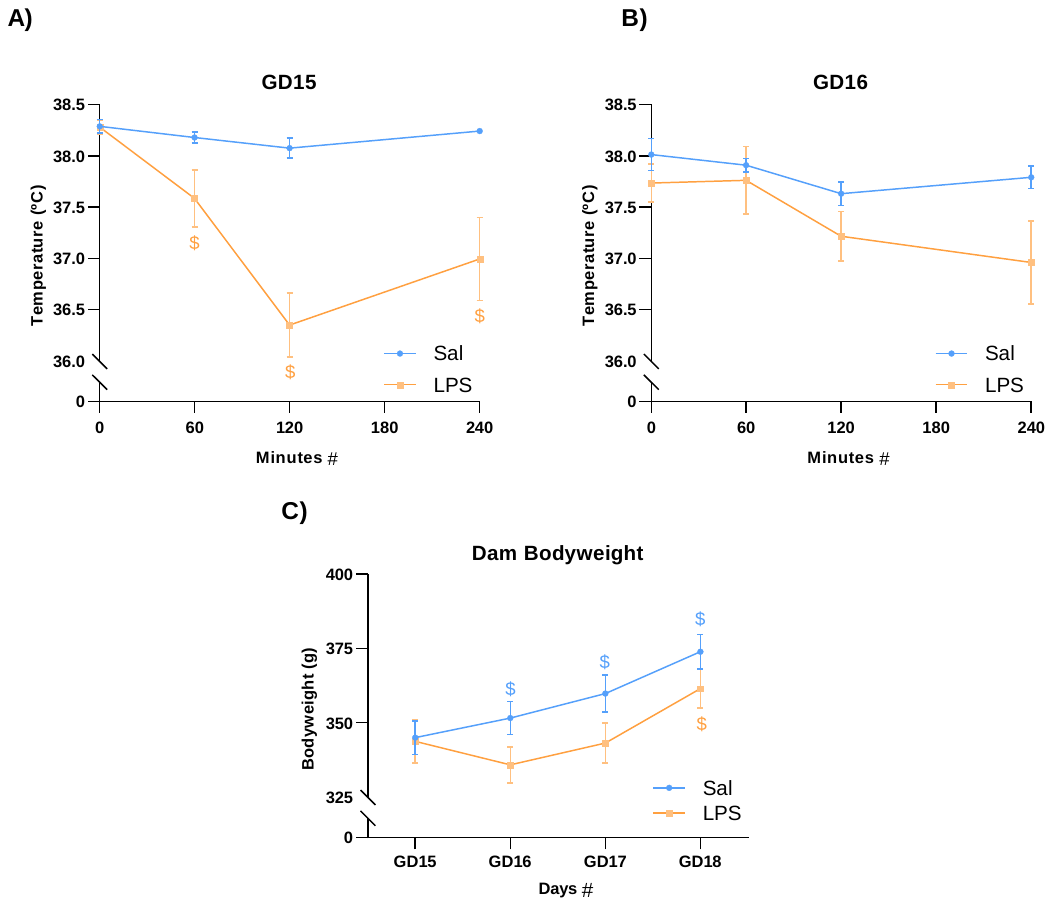

**Figure S2**: **Temperature and bodyweight changes after LPS injection**. A) and B) Rectal temperature 0, 60, 120, and 240 minutes after LPS injection on gestational days 15 and 16. C) Body weight of the mothers during the days of LPS injections and the day after the administration ended. The results are represented as mean ± SEM. n = 8 animals. *p < 0.05 between-subject effects; #p < 0.05 general within-subject effects; $ p < 0.05 differences compared to T_0_.

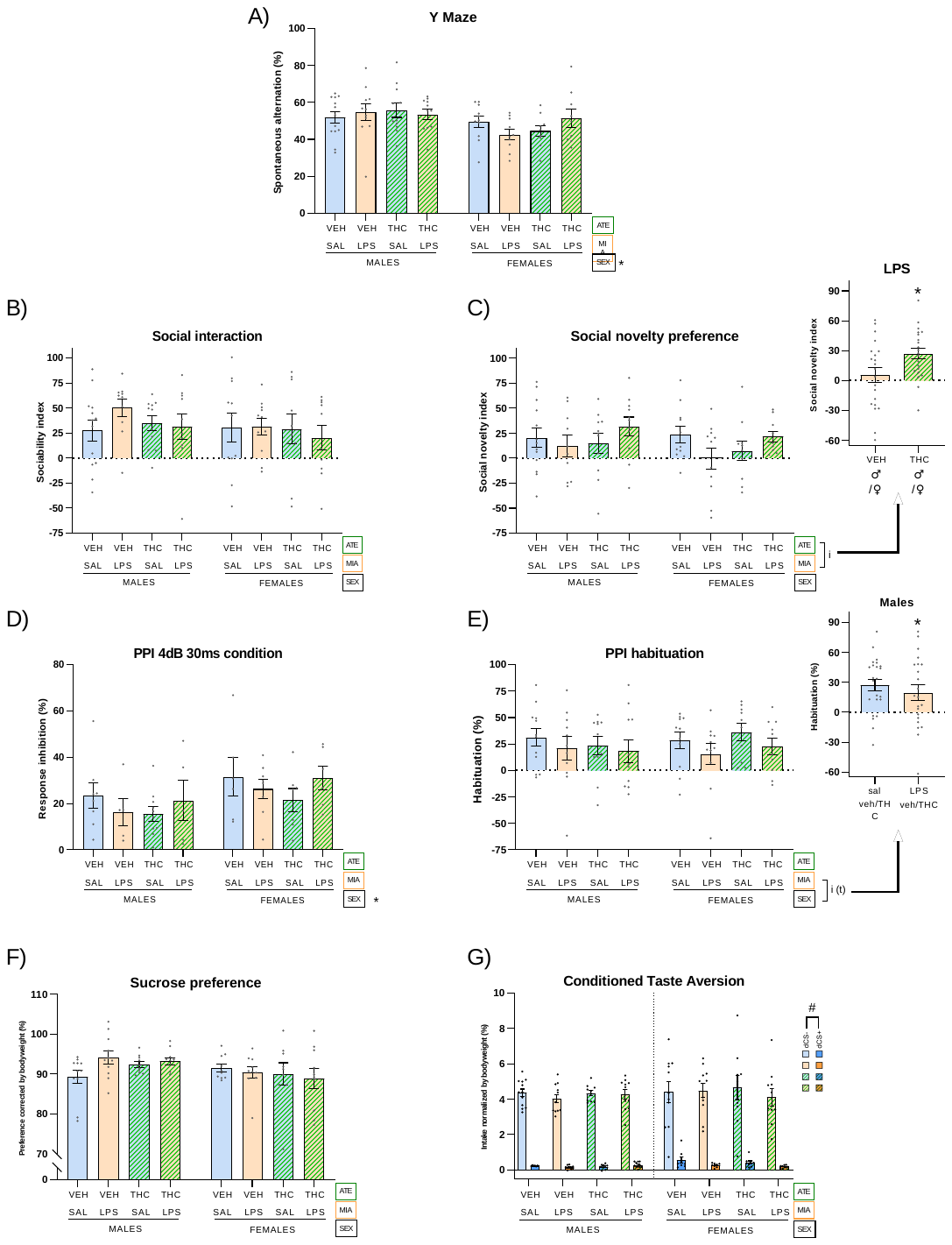

**Figure S3:** **Behavioral tests performed at adulthood to study symptom dimensions of schizophrenia**. A) Percentage of valid triads in the Y-maze relative to possible triads. A valid triad is understood as entry into 3 arms without re-entring in any of them. The number of possible triads has been calculated as the total entries minus two. B) Sociability index calculated as the time interacting with the first conspecific minus the time interacting with the inanimate object divided by the sum of both. C) Social novelty preference index calculated as the time interacting with newly introduced conspecific minus the time interacting with the familiar conspecific divided by the sum of both. D) Percentage of inhibition of the startle response by a prepulse 4 dB above the background noise and separated from the pulse (120 dB) by 30 milliseconds. E) Percentage of habituation to the main stimulus of 120 dB. F) Sucrose preference measured by a two-bottle test and expressed as a percentage. G) Fluid intake in the conditioned taste aversion taste normalised to body weight. Details of the results obtained in the statistica analysis are found in tables S1, S2 and S3. Results are represented as mean ± SEM. n in Tables R2.4. Interactions are represented with brackets and accompanied by a simplified graph. *p < 0.05 main between-subject effects; "t" (0.05 < p < 0.06) trend; # main within-subjects effect (p < 0.05). Inserts depict significant simple effects obtained after a significant interaction.

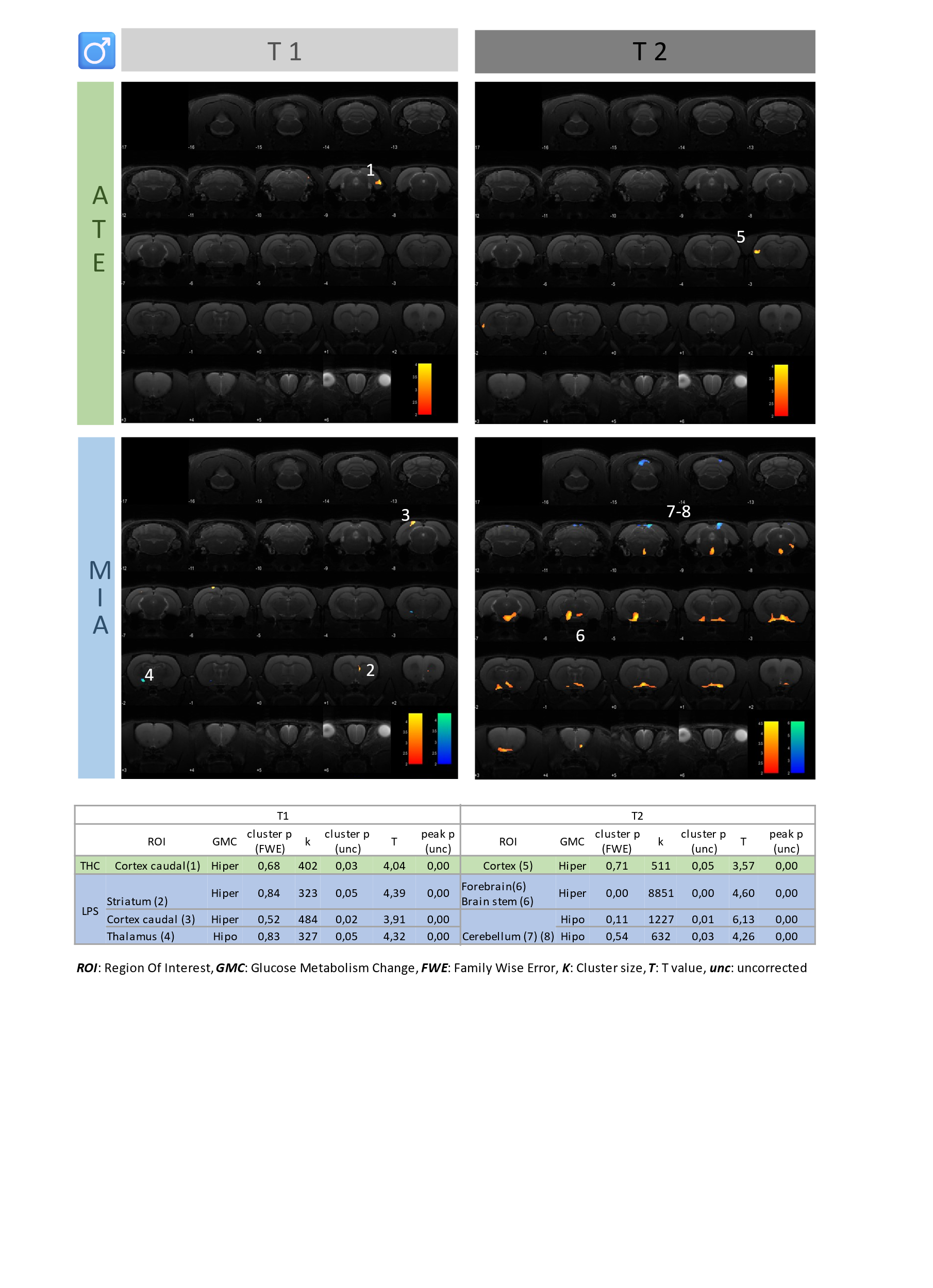
**Figure S4.** **PET analysis of brain activity in males.** A) T maps overlapped on an MR image showing changes in brain glucose metabolism in male rats. Reductions in brain activity are shown in blue and increases in red. The left images show the main effects of MIA and ATE at PND60 (T1) and the right images at PND120 (T2). T-maps were obtained as the results of a full factorial analysis with SPM12 with a p value < 0.01, uncorrected, for both factors (MIA and ATE). B) Detailed SPM12 data per time point (T1 right and T2 left) and factor (MIA bottom and ATE top) representing statistical data of each cluster. The main effect observed was the hyperactivation produced by MIA at T2 in ventral areas such as the pons and some thalamic, hypothalamic and basal forebrain nuclei.

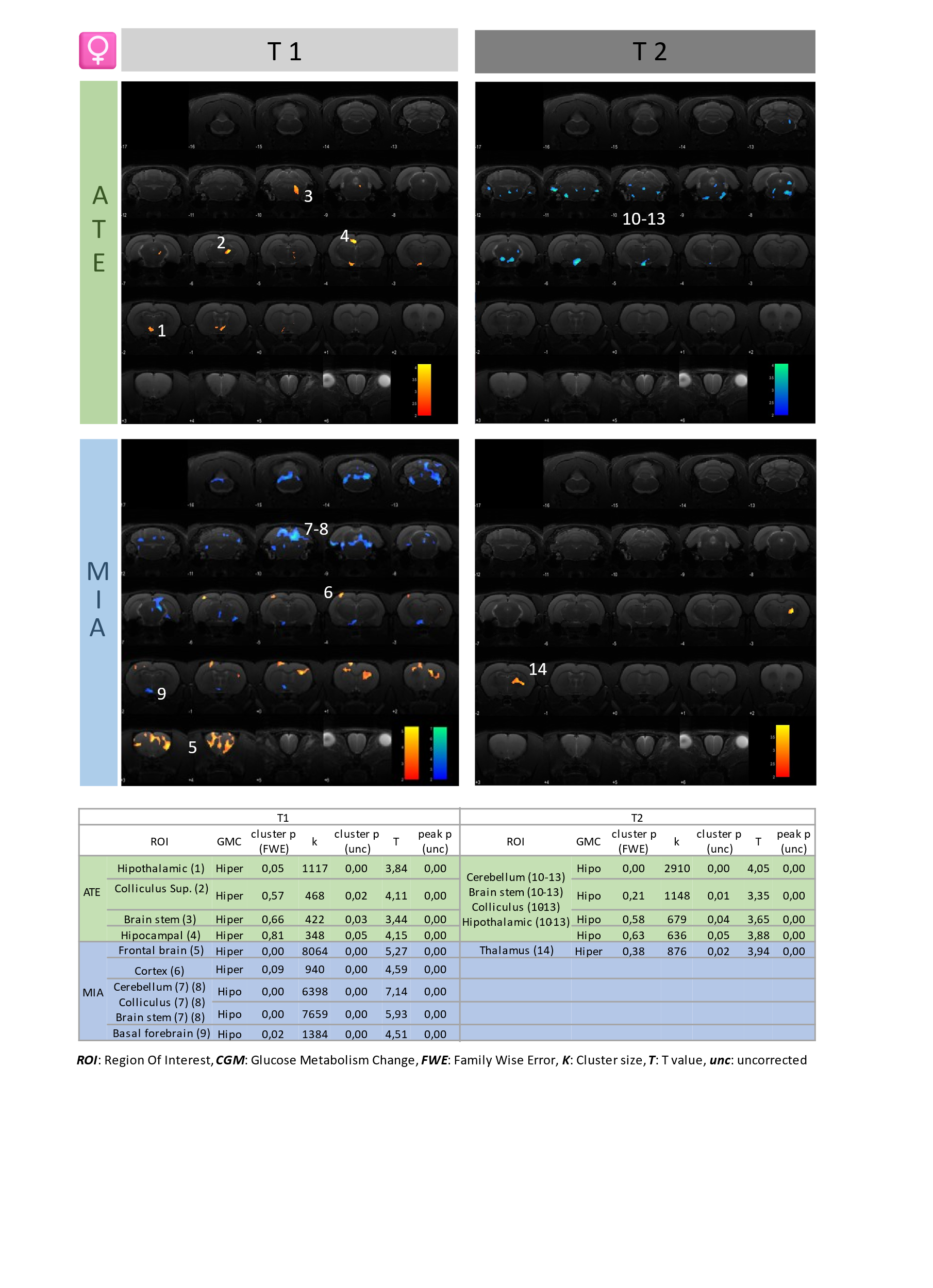

**Figure S5. PET analysis of brain activity in females.** A) T maps overlapped on an MR image showing changes in brain glucose metabolism in female rats. Reductions in brain activity are shown in blue and increases in red. The left images represent the main effects of MIA and ATE at PND60 (T1) and right images at PND120 (T2). T-maps were obtained as the results of a full factorial analysis with SPM12 at p-value < 0.01 uncorrected for both factors (MIA and ATE). B) Detailed SPM12 data per time point (T1 right and T2 left) and factor (MIA bottom and ATE top) representing statistical data of each cluster. The effect of ATE in T1 showed several clusters of hyperactivation in the pons, thalamus, cortex and basal forebrain which disappeared at T2, when a pattern of hypoactivation in the anterior regions of the brainstem, cerebellum, colliculus and thalamic/hypothalamic regions emerged. MIA at T1 produced both, hyper- and hypoactivation clusters in the forebrain and hindbrain areas, respectively, which were not present at T2.

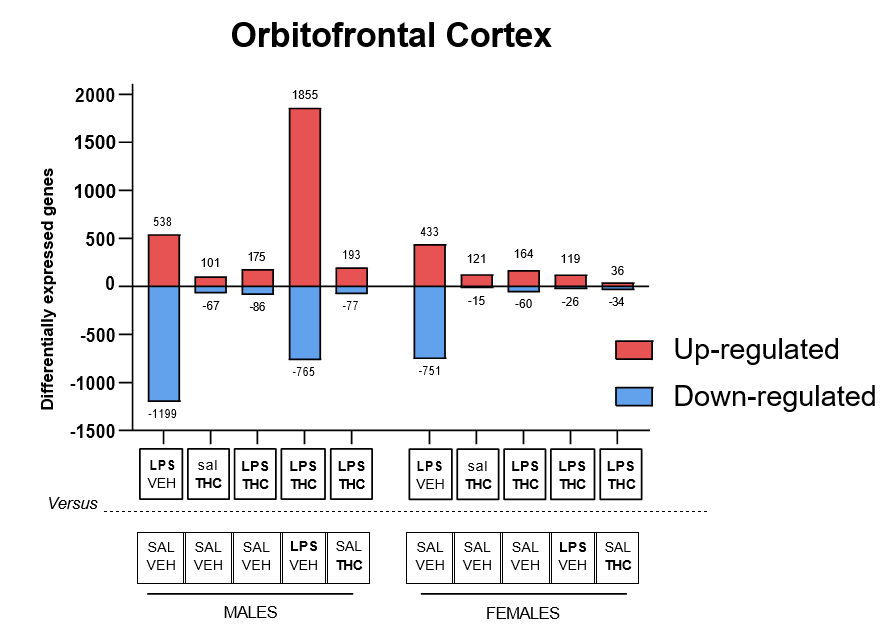

**Figure S6: Number of differentially expressed genes in OFC.** The number of differentially expressed genes in each of the comparisons is represented.

B

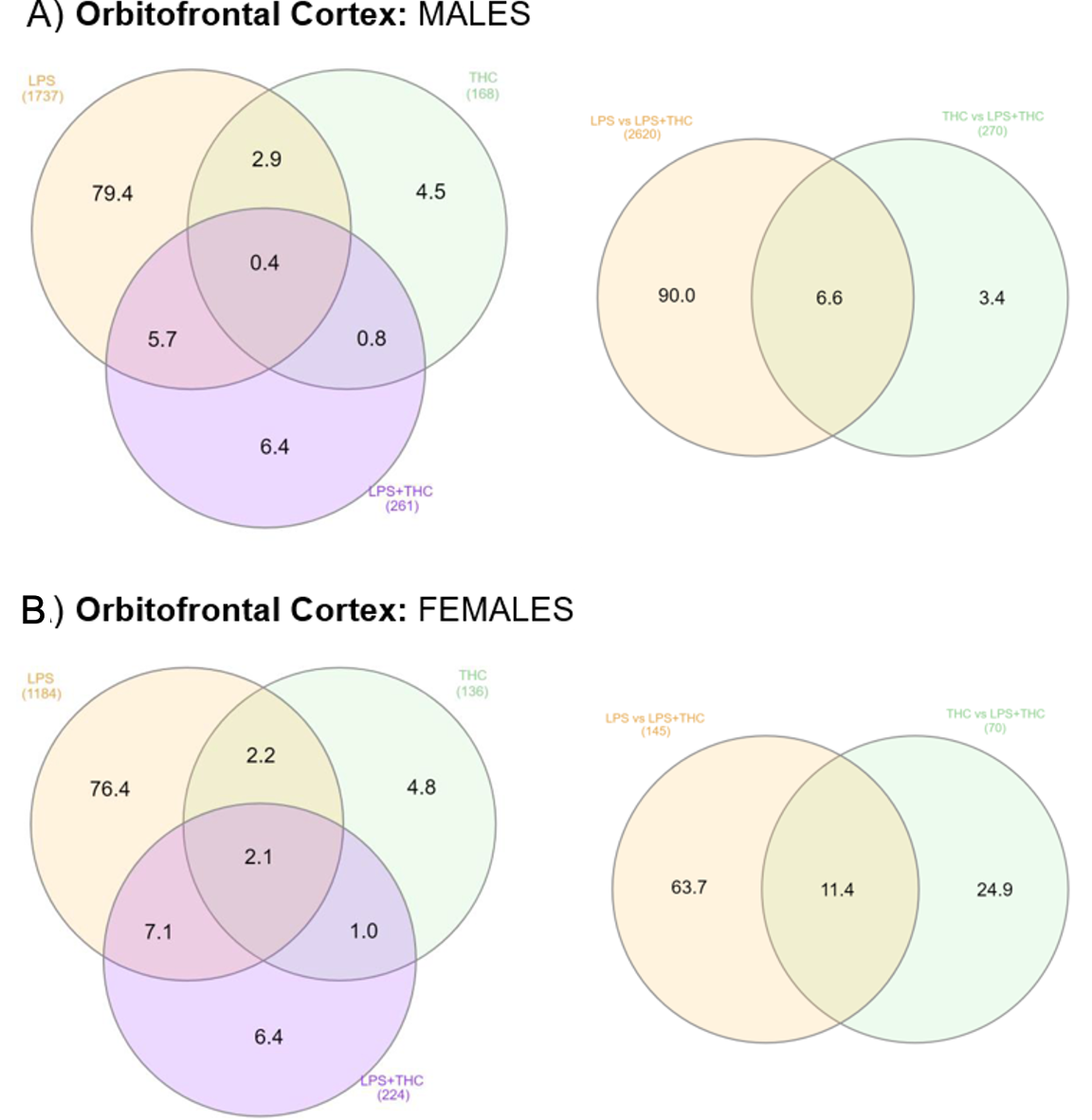

**Figure S7:** **Distribution of differentially expressed genes in the OFC.** A) and B) Left: Percentages of differentially DEGs are represented in a Venn diagram comparing the experimental groups LPS+veh (yellow), saline+THC (green), and LPS+THC (purple) with saline+vehicle. A) and B) Right: The distribution of DEGs is shown comparing the LPS+THC group with LPS+vehicle (yellow) and saline+THC (green). The total number of DEGs is shown in brackets. Created with InteractiVenn (Heberle et al., 2015).

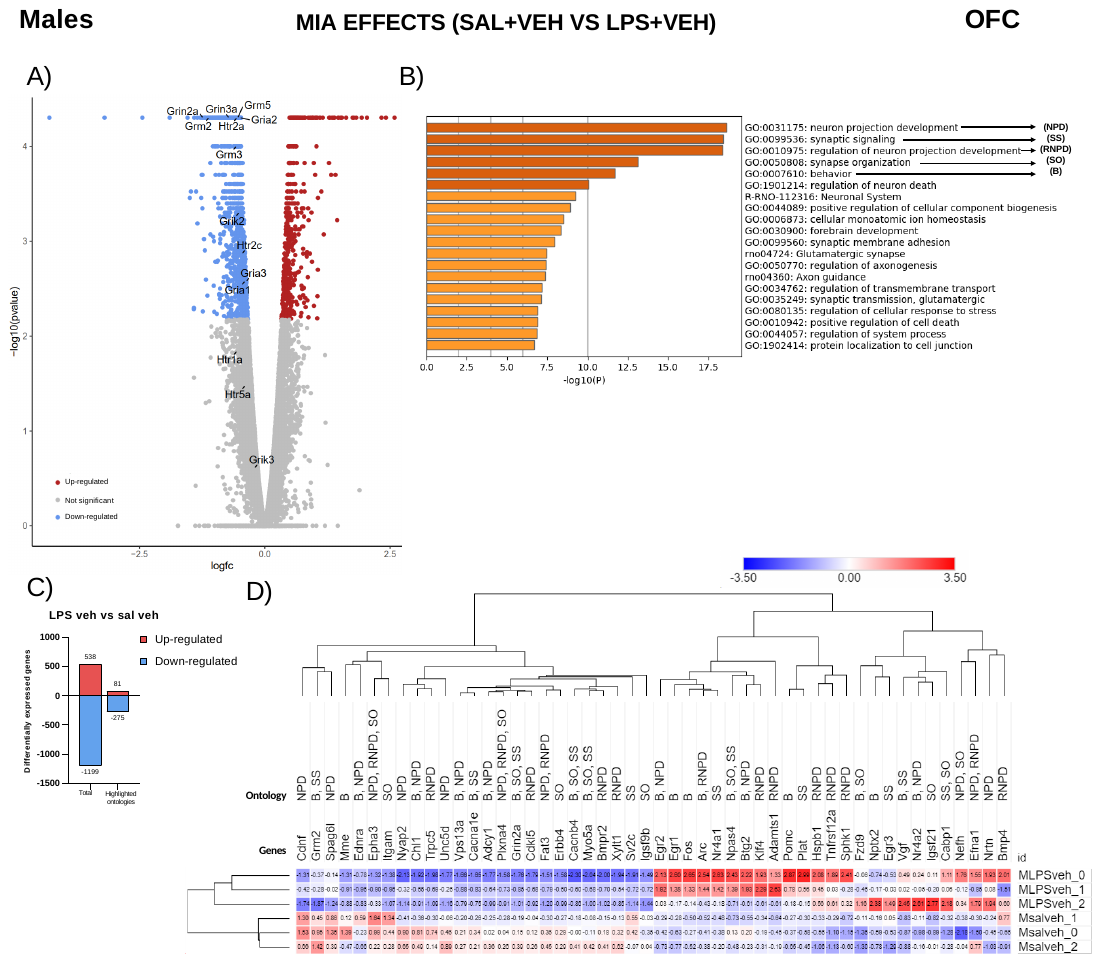

**Figure S8: MIA effects in OFC transcriptome in males**. A) -Log10 of the unadjusted p-value is plotted against the logarithm of FC (fold change). Genes differentially expressed upregulated are shown in red, while those downregulated are shown in blue (adjusted p-value < 0.05). Genes of glutamatergic and serotoninergic receptors, or their subunits, affected by exposure to MIA, ATE, or their combination are depicted. B) Ontologies significantly enriched with differentially expressed genes. C) Graphical representation of the number of differentially expressed genes, both total (Total) and only from the 5 highlighted ontologies (highlighted ontologies). D) Heatmap showing the number of fragments from each sample normalized to Z-score using the mean and standard deviation of fragments from all males. All genes belonging to the highlighted ontologies are represented.

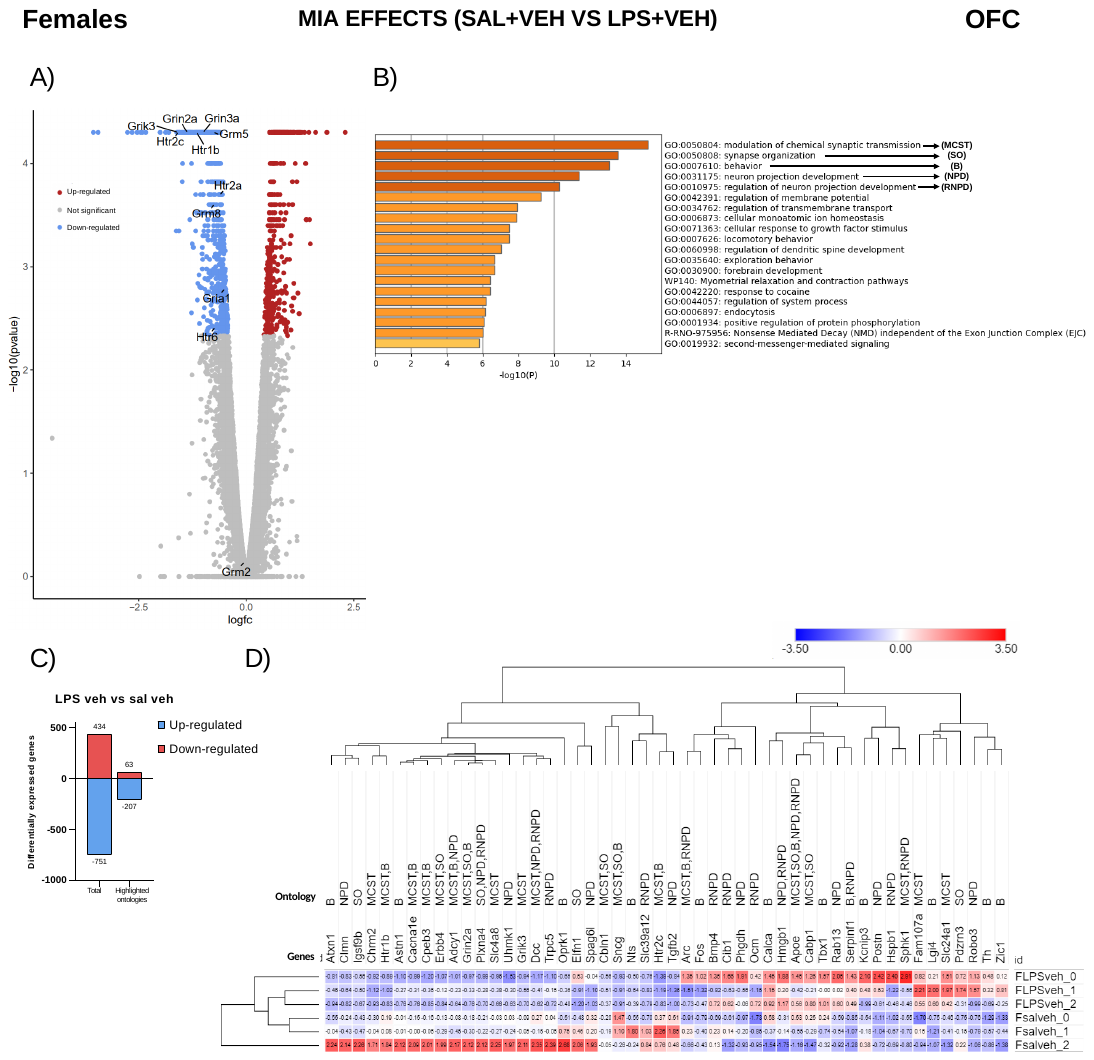

**Figure S9: MIA effects in OFC transcriptome in females**. A) -Log10 of the unadjusted p-value is plotted against the logarithm of FC (fold change). Genes differentially expressed upregulated are shown in red, while those downregulated are shown in blue (adjusted p-value < 0.05). Genes of glutamatergic and serotoninergic receptors, or their subunits, affected by exposure to MIA, ATE, or their combination are depicted. B) Ontologies significantly enriched with differentially expressed genes. C) Graphical representation of the number of differentially expressed genes, both total (Total) and only from the 5 highlighted ontologies (highlighted ontologies). D) Heatmap showing the number of fragments from each sample normalized to Z-score using the mean and standard deviation of fragments from all males. All genes belonging to the highlighted ontologies are represented.

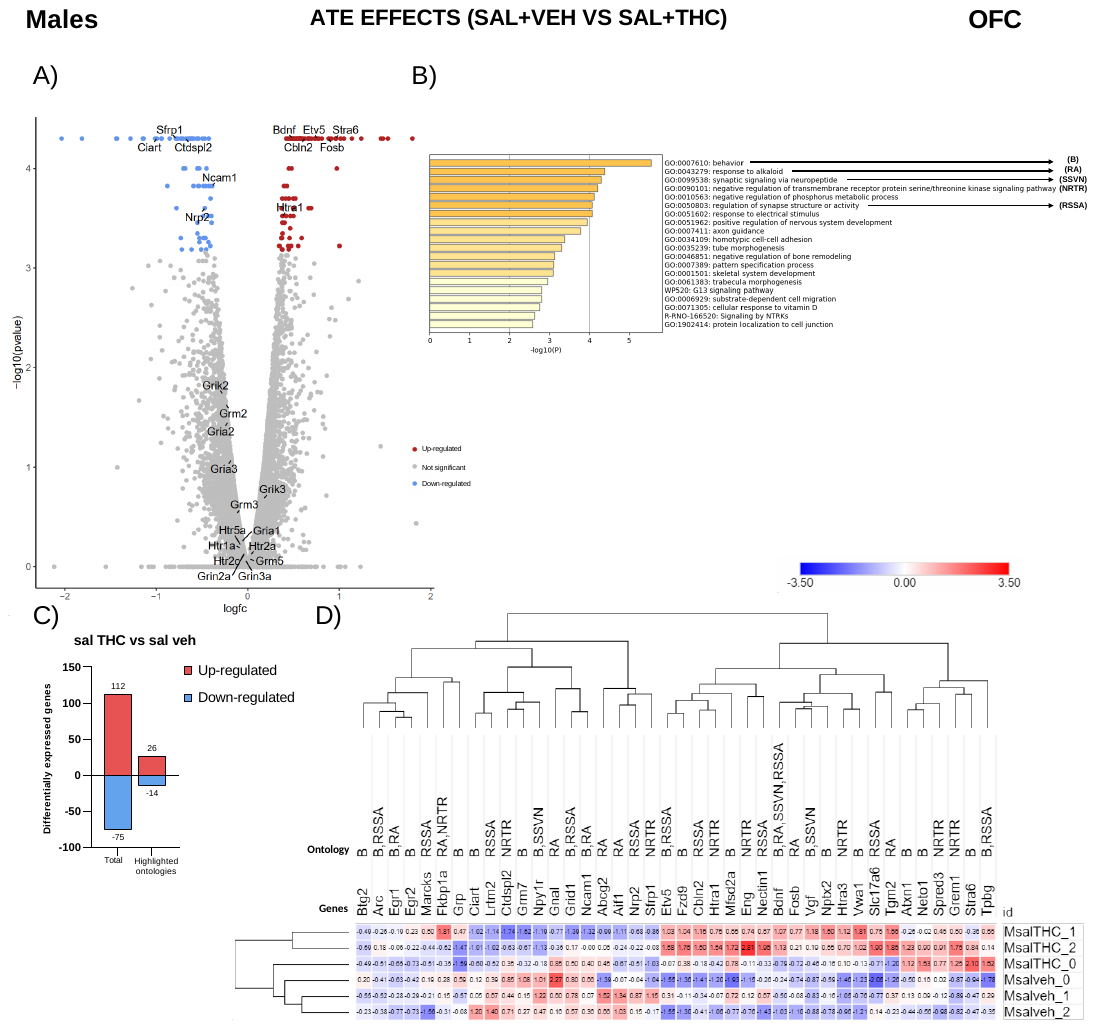

**Figure S10: ATE effects in OFC transcriptome in males**. A) -Log10 of the unadjusted p-value is plotted against the logarithm of FC (fold change). Genes differentially expressed upregulated are shown in red, while those downregulated are shown in blue (adjusted p-value < 0.05). Genes of glutamatergic and serotoninergic receptors, or their subunits, affected by exposure to MIA, ATE, or their combination are depicted. B) Ontologies significantly enriched with differentially expressed genes. C) Graphical representation of the number of differentially expressed genes, both total (Total) and only from the 5 highlighted ontologies (highlighted ontologies). D) Heatmap showing the number of fragments from each sample normalized to Z-score using the mean and standard deviation of fragments from all males. All genes belonging to the highlighted ontologies are represented.

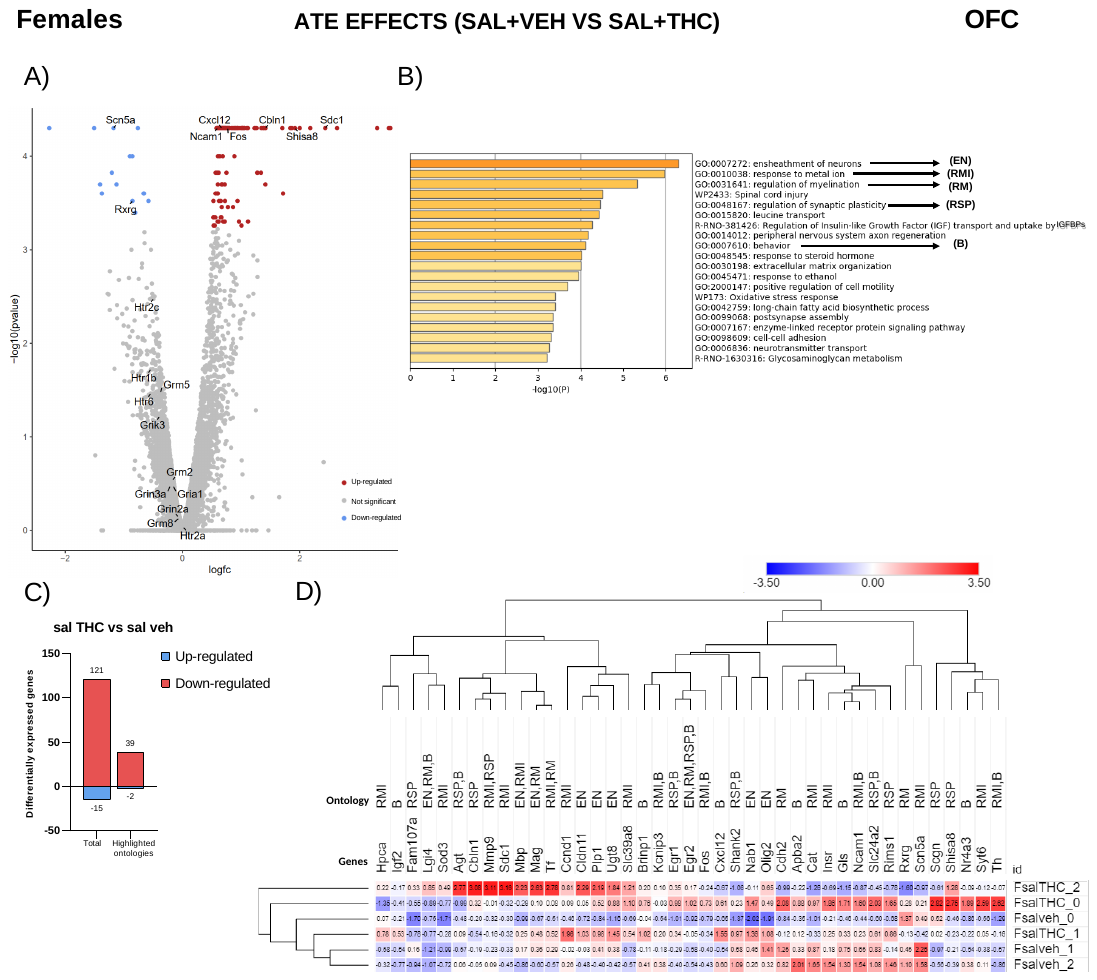

**Figure S11: ATE effects in OFC transcriptome in females**. A) -Log10 of the unadjusted p-value is plotted against the logarithm of FC (fold change). Genes differentially expressed upregulated are shown in red, while those downregulated are shown in blue (adjusted p-value < 0.05). Genes of glutamatergic and serotoninergic receptors, or their subunits, affected by exposure to MIA, ATE, or their combination are depicted. B) Ontologies significantly enriched with differentially expressed genes. C) Graphical representation of the number of differentially expressed genes, both total (Total) and only from the 5 highlighted ontologies (highlighted ontologies). D) Heatmap showing the number of fragments from each sample normalized to Z-score using the mean and standard deviation of fragments from all males. All genes belonging to the highlighted ontologies are represented.

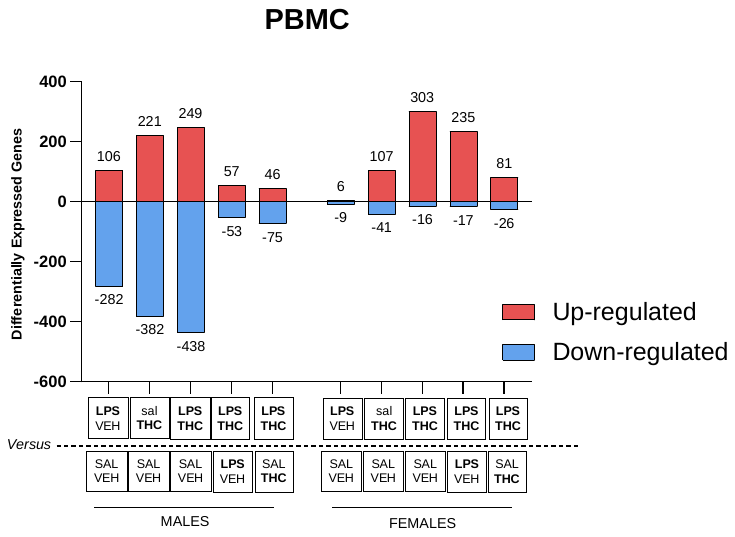

**Figure S12:** **Number of differentially expressed genes in the PMBCs.** The number of differentially expressed genes in each of the comparisons is represented.

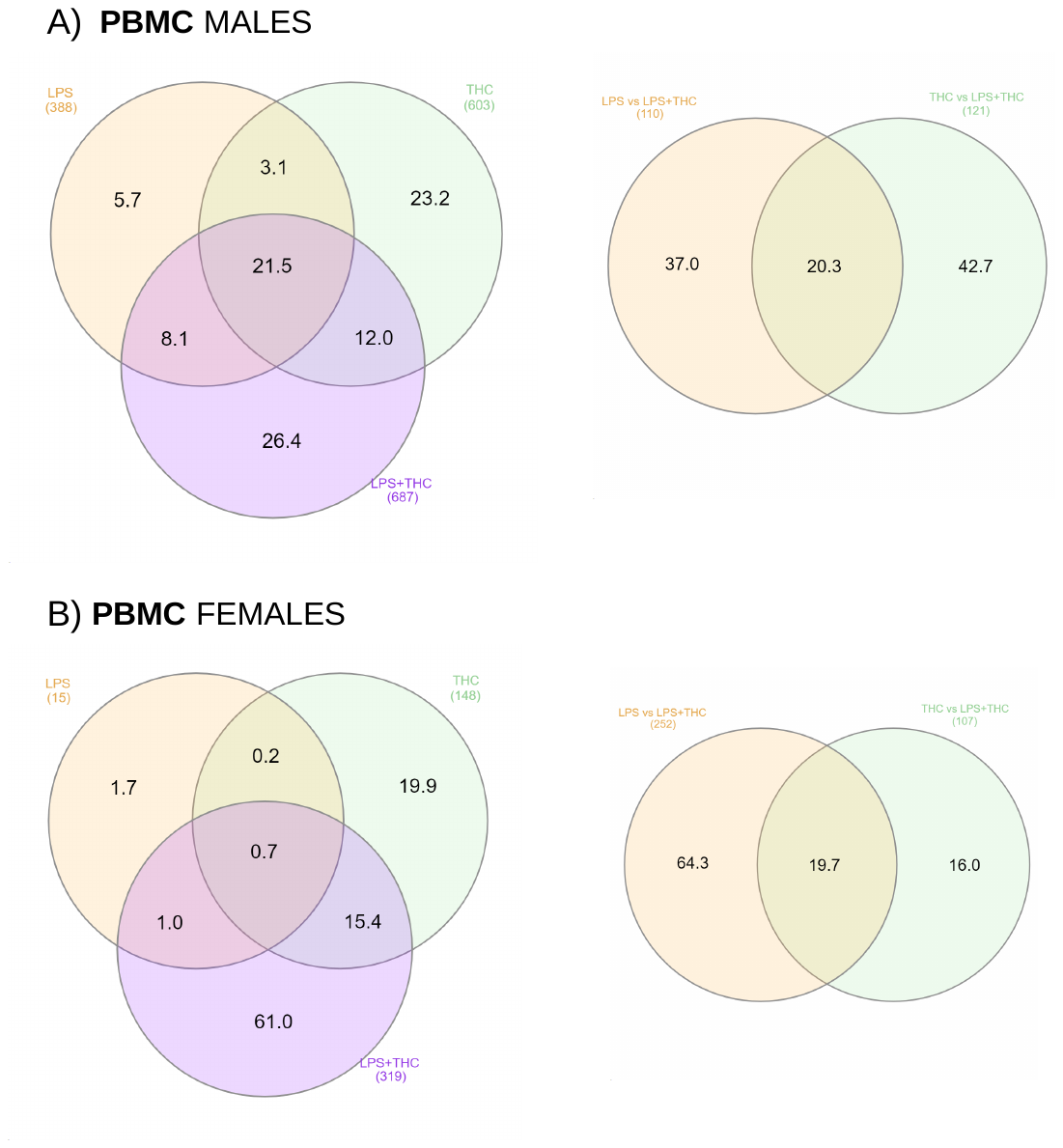

**Figure S13:** **Distribution of differentially expressed genes in the PBMCs.** A) and B) Left: Percentages of differentially DEGs are represented in a Venn diagram comparing the experimental groups LPS+vehicle (yellow), saline+THC (green), and LPS+THC (purple) with saline+vehicle. A) and B) Right: The distribution of DEGs is shown comparing the LPS+THC group with LPS+vehicle (yellow) and saline+THC (green). The total number of DEGs is shown in brackets. Created with InteractiVenn (Heberle et al., 2015).

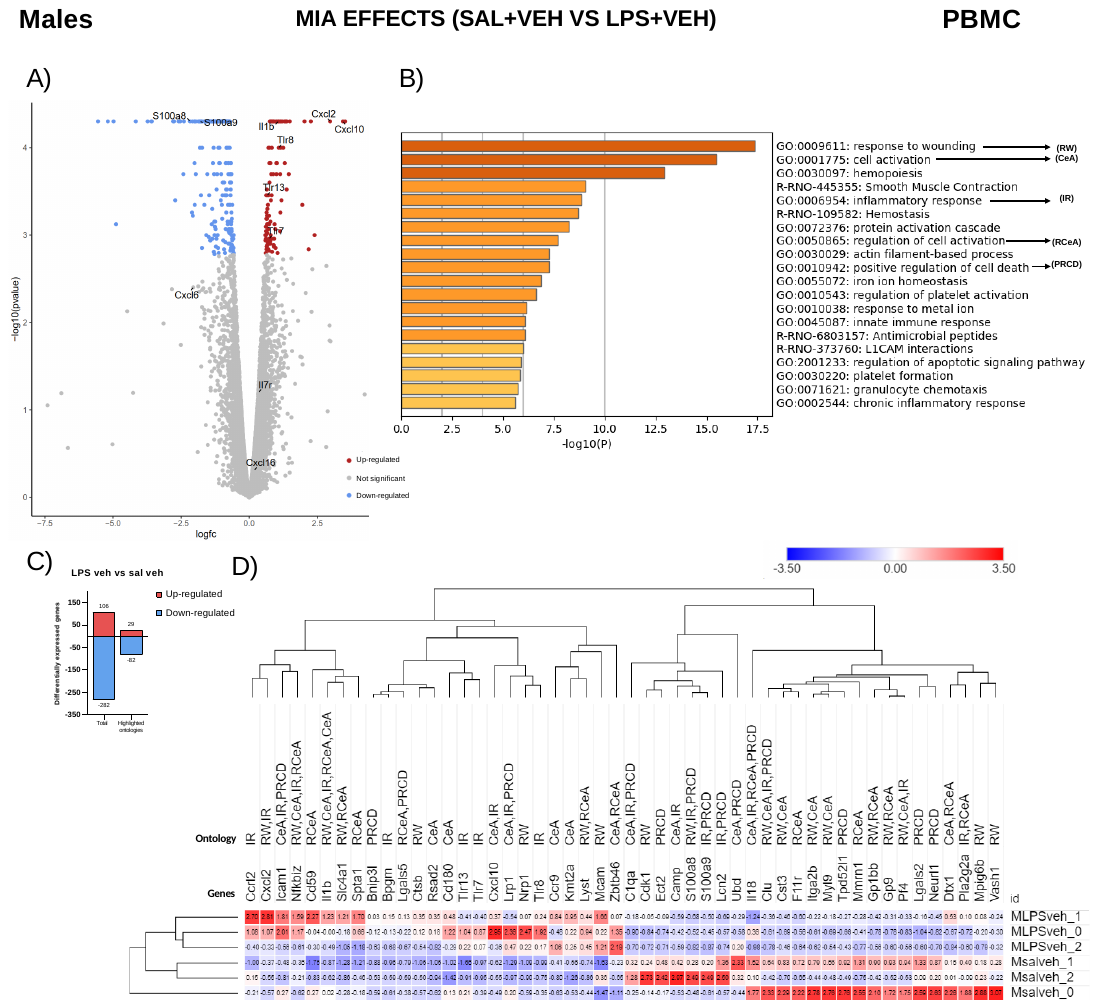

**Figure S14: MIA effects in PBMC transcriptome in males**. A) -Log10 of the unadjusted p-value is plotted against the logarithm of FC (fold change). Genes differentially expressed upregulated are shown in red, while those downregulated are shown in blue (adjusted p-value < 0.05). Genes of glutamatergic and serotoninergic receptors, or their subunits, affected by exposure to MIA, ATE, or their combination are depicted. B) Ontologies significantly enriched with differentially expressed genes. C) Graphical representation of the number of differentially expressed genes, both total (Total) and only from the 5 highlighted ontologies (highlighted ontologies). D) Heatmap showing the number of fragments from each sample normalized to Z-score using the mean and standard deviation of fragments from all males. All genes belonging to the highlighted ontologies are represented.

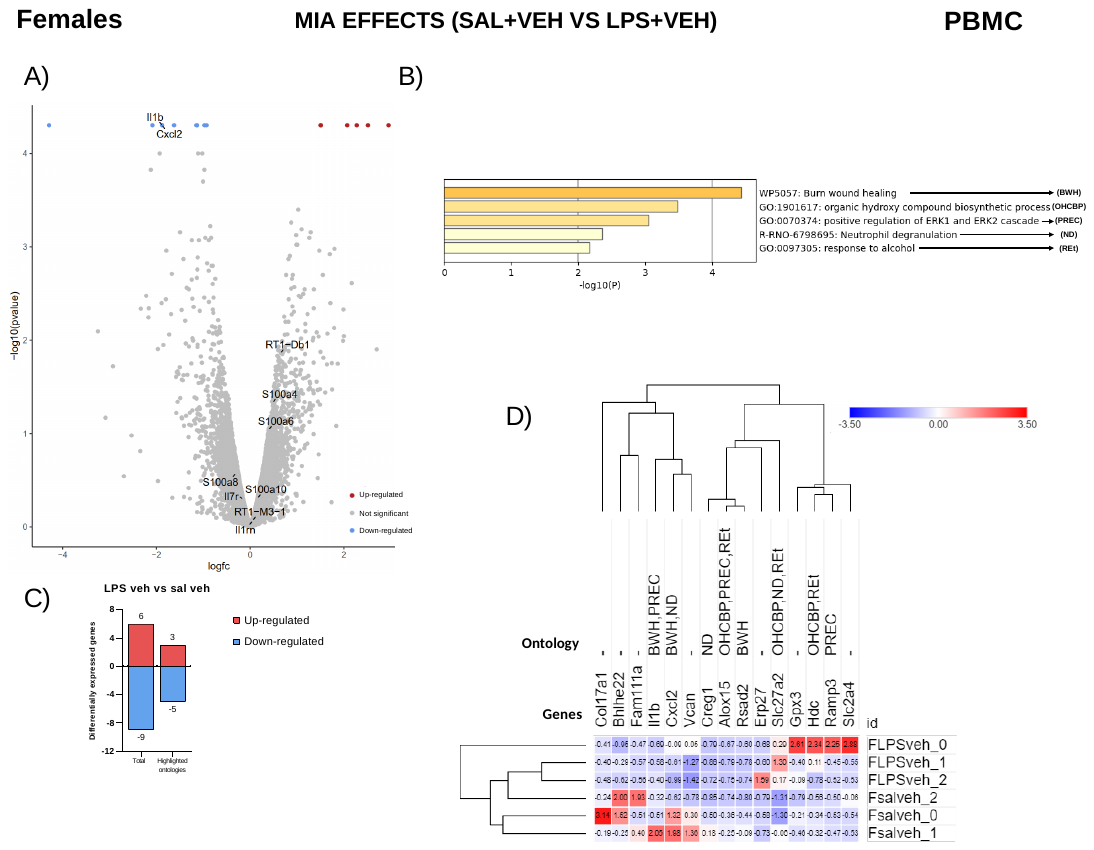

**Figure S15: MIA effects in PBMC transcriptome in females**. A) -Log10 of the unadjusted p-value is plotted against the logarithm of FC (fold change). Genes differentially expressed upregulated are shown in red, while those downregulated are shown in blue (adjusted p-value < 0.05). Genes of glutamatergic and serotoninergic receptors, or their subunits, affected by exposure to MIA, ATE, or their combination are depicted. B) Ontologies significantly enriched with differentially expressed genes. C) Graphical representation of the number of differentially expressed genes, both total (Total) and only from the 5 highlighted ontologies (highlighted ontologies). D) Heatmap showing the number of fragments from each sample normalized to Z-score using the mean and standard deviation of fragments from all males. All genes belonging to the highlighted ontologies are represented.

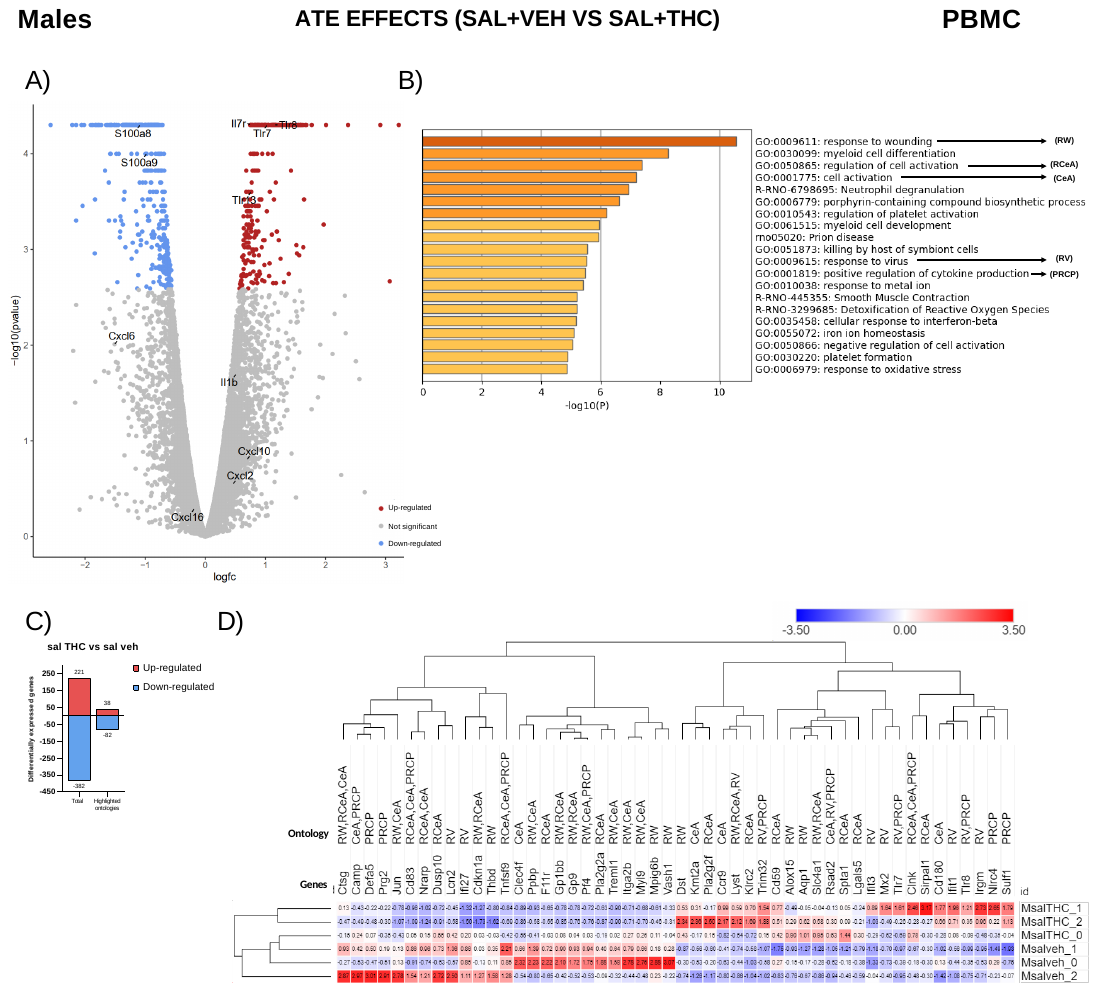

**Figure S16: ATE effects in PBMC transcriptome in males**. A) -Log10 of the unadjusted p-value is plotted against the logarithm of FC (fold change). Genes differentially expressed upregulated are shown in red, while those downregulated are shown in blue (adjusted p-value < 0.05). Genes of glutamatergic and serotoninergic receptors, or their subunits, affected by exposure to MIA, ATE, or their combination are depicted. B) Ontologies significantly enriched with differentially expressed genes. C) Graphical representation of the number of differentially expressed genes, both total (Total) and only from the 5 highlighted ontologies (highlighted ontologies). D) Heatmap showing the number of fragments from each sample normalized to Z-score using the mean and standard deviation of fragments from all males. All genes belonging to the highlighted ontologies are represented.

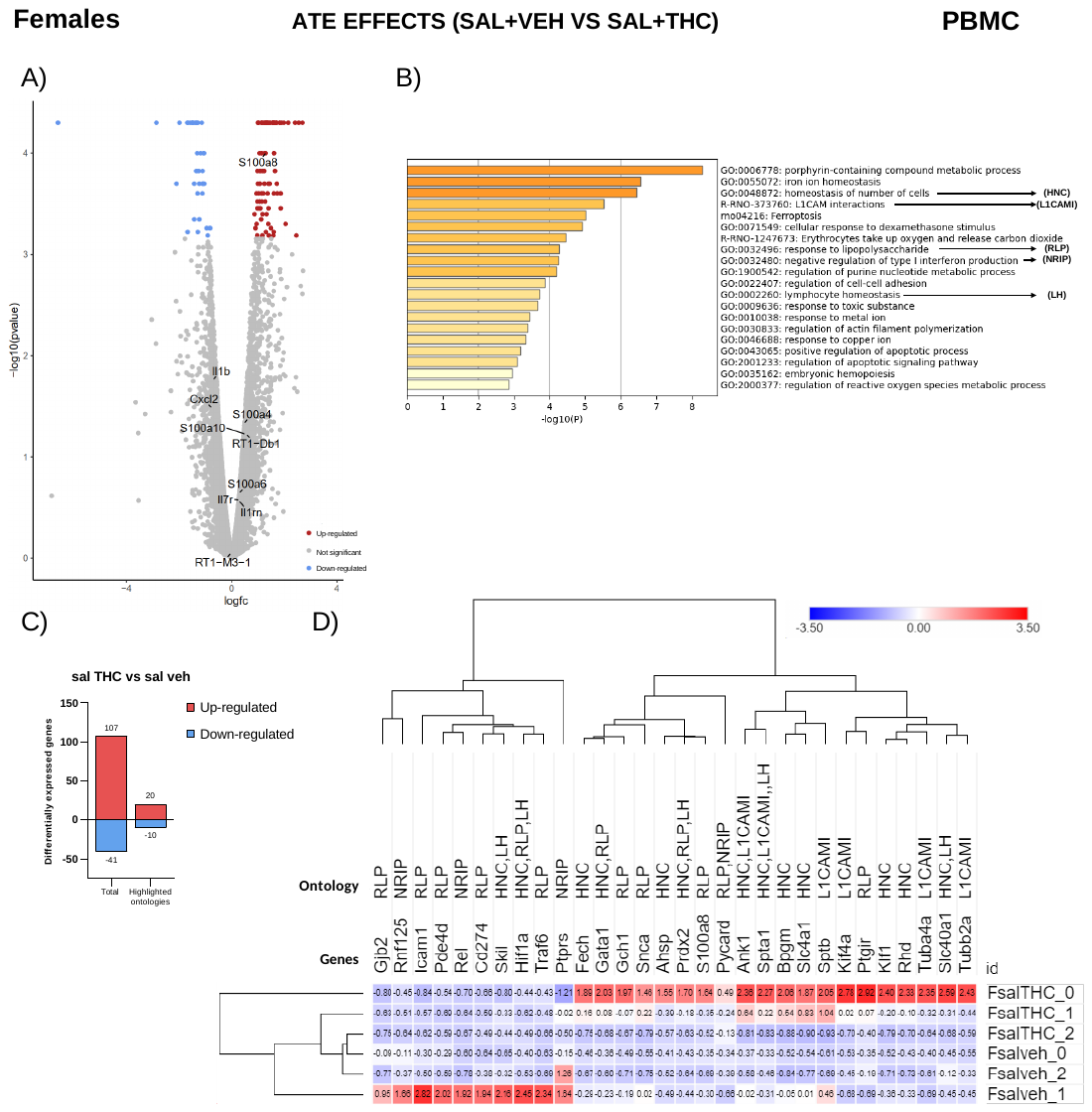

**Figure S17: ATE effects in PBMC transcriptome in males**. A) -Log10 of the unadjusted p-value is plotted against the logarithm of FC (fold change). Genes differentially expressed upregulated are shown in red, while those downregulated are shown in blue (adjusted p-value < 0.05). Genes of glutamatergic and serotoninergic receptors, or their subunits, affected by exposure to MIA, ATE, or their combination are depicted. B) Ontologies significantly enriched with differentially expressed genes. C) Graphical representation of the number of differentially expressed genes, both total (Total) and only from the 5 highlighted ontologies (highlighted ontologies). D) Heatmap showing the number of fragments from each sample normalized to Z-score using the mean and standard deviation of fragments from all males. All genes belonging to the highlighted ontologies are represented.
